# Context-dependent and dynamic functional influence of corticothalamic pathways to first- and higher-order visual thalamus

**DOI:** 10.1101/738203

**Authors:** Megan A. Kirchgessner, Alexis D. Franklin, Edward M. Callaway

## Abstract

Layer 6 (L6) is the sole purveyor of corticothalamic (CT) feedback to first-order thalamus and also sends projections to higher-order thalamus, yet how it engages the full corticothalamic circuit to contribute to sensory processing in an awake animal remains unknown. We sought to elucidate the functional impact of L6CT projections from primary visual cortex to visual thalamic nuclei dLGN (first-order) and pulvinar (higher-order) using optogenetics and extracellular electrophysiology in awake mice. While sustained L6CT photostimulation suppresses activity in both visual thalamic nuclei *in vivo*, moderate-frequency (10Hz) stimulation powerfully facilitates thalamic spiking. We show that each stimulation paradigm differentially influences the balance between monosynaptic excitatory and disynaptic inhibitory corticothalamic pathways to dLGN and pulvinar as well as the prevalence of burst versus tonic firing. Altogether, our results support a model in which L6CTs modulate first- and higher-order thalamus through parallel excitatory and inhibitory pathways that are highly dynamic and context-dependent.

**Significance:** Layer 6 corticothalamic (L6CT) projections play important modulatory roles in thalamic processing, yet *how* this modulation is executed is unclear. While some studies suggest fundamentally inhibitory influence of L6CTs over first-order thalamus, potential complex, frequency-dependent effects have not been investigated *in vivo.* Moreover, how L6CTs affect higher-order nuclei *in vivo* has not been explored. This study utilizes various optogenetic manipulations of L6CTs with single-unit recordings from multiple thalamic nuclei in awake mice to address these questions. Our results illustrate similar effects of L6CTs on first- and higher-order visual thalamic nuclei, yet very different effects within-nucleus depending on how L6CTs are engaged. These findings suggest that L6CT modulation is not simply inhibitory by nature, but instead is dynamic and context-dependent.

## Introduction

While the flow of information from thalamus to cortex is widely appreciated as a critical step in sensory processing, the significance of a given cortical area’s projection back to thalamus is considerably less clear. In the case of first-order nuclei, like the dorsal lateral geniculate nucleus (dLGN) in the visual system, this corticothalamic feedback originates from layer 6. Layer 6 corticothalamic neurons (L6CTs) have been classically described as providing “modulatory” feedback to dLGN (1) that may influence response gain (2–6), temporal precision (7, 8), spatiotemporal filtering (5, 8, 9), sensory adaptation (10), and firing mode (6, 9, 10). Still, how these L6CTs might perform these various functions is not well understood. Moreover, many L6CTs also project to higher-order thalamus, such as the visual pulvinar (also known as the lateral posterior nucleus, or LP, in rodents) (11). Extensive anatomical and physiological evidence demonstrates similar “modulatory” characteristics of L6CT projections to both first- and higher-order thalamus (12–14), yet this pathway has not been investigated *in vivo*. Thus, many questions remain with regards to the nature of corticothalamic feedback as a general feature of sensory circuits and whether these same principles hold across different classes of thalamic nuclei.

One such question is how L6CTs influence their thalamic targets during sensory processing in an awake animal. On one hand, previous observations of dramatically reduced visual responses recorded in dLGN of anesthetized mice during V1 L6CT optogenetic activation (3, 4) suggest that L6CT feedback may be fundamentally inhibitory, likely through a disynaptic inhibitory pathway through the GABAergic thalamic reticular nucleus (TRN). However, other studies in the visual as well as other sensory systems disagree, finding no change or even increased activity in first-order thalamus with L6CT photoactivation (8, 15, 16). There are many challenges in interpreting these conflicting findings. First, the great majority of these studies were conducted under anesthesia, which has been shown to affect both spontaneous and visually-evoked firing rates in dLGN (17). Second, most of these prior studies have in common that they delivered continuous light for photostimulation, with which different levels of opsin expression as well as duration and intensity of light stimulation might yield different effects due to the uncontrolled nature of the manipulation. Finally, optogenetically inactivating L6CTs has mixed effects in dLGN (4), suggesting that their natural function is not to invariably suppress their thalamic targets.

An alternative explanation could be that the level and manner of L6CTs’ activation may determine how they influence their thalamic targets. For instance, the effects of L6CT optogenetic stimulation on first-order thalamus VPm in the somatosensory *in vitro* slice preparation have been shown to “switch” from being net-suppressing to net-facilitating with higher-frequency (10Hz) L6CT stimulation (18). This frequency-dependence has been explained by the different short-term plasticity characteristics at different synapses in the full corticothalamic circuit, since the competing monosynaptic excitatory and disynaptic inhibitory (via TRN) routes to first-order thalamus are net-facilitating and net-depressing, respectively (18). Previous studies have not used controlled L6CT optogenetic manipulations in awake animals or probed *in vivo* L6CT effects on a higher-order thalamic nucleus. Therefore, it remains to be seen whether L6CT projections can exert flexible, bi-directional influence on thalamic activity in the visual system, in different classes of thalamic nuclei, and *in vivo*.

To address these questions, we have recorded extracellular single-unit activity from dLGN, pulvinar, and TRN in awake mice. We optogenetically manipulated L6CTs in primary visual cortex with both controlled (photostimulation trains) and uncontrolled (continuous light) methods for photostimulation. While we observe similar influences of L6CTs on dLGN and pulvinar, different photostimulation conditions had strikingly different effects on thalamic firing rates, firing mode, and the balance with activity changes in the TRN. Our results thus provide novel evidence that L6CTs are capable of dynamically influencing activity in both their first-and higher-order thalamic targets.

## Results

### Sustained L6CT photostimulation suppresses activity in dLGN and pulvinar *in vivo*

Before turning to more controlled stimulation of L6CT neurons, we first tested whether previous effects observed in dLGN of the anesthetized animal during sustained L6CT photostimulation are also observed in *awake* mice. Additionally, since L6CTs are hypothesized to play similar functional roles in both first- and higher-order thalamic nuclei (1), we wondered whether effects observed in dLGN would also extend to the pulvinar. To address these questions, an AAV encoding Cre-dependent ChR2-eYFP was injected into V1 of Ntsr1-Cre GN220 transgenic mice (Methods). Consistent with prior reports of the specificity of the Ntsr1-Cre transgenic line (19, 20), expression of the ChR2-eYFP fusion protein was specific to V1 layer 6 (Fig. 1B). Corticothalamic axons expressing ChR2-eYFP were readily apparent in both dLGN and pulvinar (Figs. 1C and S1A), demonstrating that L6CTs labeled by the Ntsr1-Cre line project to both visual thalamic nuclei.

**Fig. 1.**
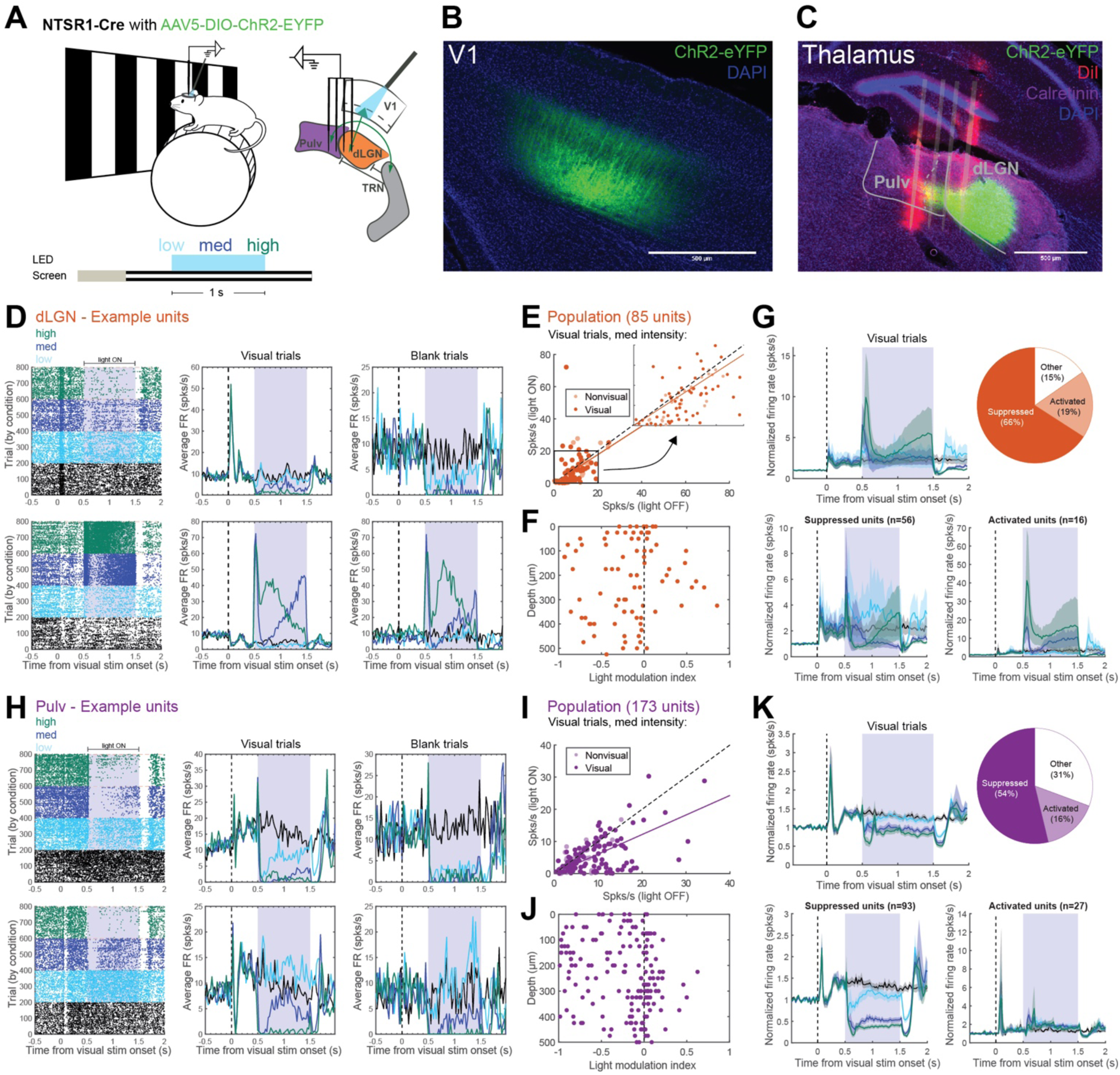
L6CT photostimulation with continuous light delivery suppresses activity in dLGN and pulvinar *in vivo*. (A) Experimental design. Left: diagram of the experimental setup and trial structure for visual and LED stimulation. Right: schematic of the L6 corticothalamic circuit, indicating recording and LED stimulation locations. (B) Coronal section depicting ChR2-eYFP cell body expression in V1 L6 and apical dendrites in L4 of a Ntsr1-Cre mouse injected with AAV5-DIO-ChR2-eYFP. (C) Coronal section of the visual thalamus, depicting ChR2-eYFP-positive axon terminals in dLGN and pulvinar. Recording tracks from a four-shank probe are labeled with DiI (red). Immunohistochemical staining for calretinin (purple) provides borders from lateral pulvinar to dLGN and medial pulvinar. (D) Two example dLGN units. Left panels: raster plots with trials organized by LED condition (trials with different conditions were interspersed during the actual experiment). The shaded area indicates photostimulation period (0.5-1.5s following visual stimulus onset). Middle panels: peristimulus time histograms (PSTHs) of average firing rates during visual trials for each condition, shown over time relative to visual stimulus onset. Right panels: PSTHs of average firing rates during blank trials (grey screen). (E) Average firing rates during the 1-second photostimulation period from visual trials, with versus without medium-intensity L6CT photostimulation. Inset: expanded scatter plot from area within the square (0-20 spks/s). Saturated points indicate visually-responsive units. (F) Light modulation index (<0 suppressed, >0 activated) by depth (distance from highest channel on the probe with a visually-responsive unit). (G) Effects of different LED intensities on units’ visually-evoked activity in dLGN. Top, left: mean normalized PSTH (normalized to each unit’s prestimulus firing rate) across all dLGN units. Top, right: proportion of units which were significantly suppressed or activated in two or more conditions (units not passing this criteria considered “other”). Bottom: average normalized PSTH across suppressed (left) and activated (right) units. Shading indicates ±1 standard error of the mean. (H-K) Same as (D-G) but for units recorded in lateral pulvinar.

Single-unit activity was recorded in the visual thalamus of awake, head-fixed mice using high-density, multi-shank microelectrode arrays (21). Probes were coated with lipophilic dye (DiI) in order to visualize electrode tracks and determine which thalamic nucleus was sampled by each shank (Fig. 1C). Since the mouse pulvinar is not uniformly innervated by V1 (22–24), calretinin expression was used to distinguish between pulvinar subdivisions (Fig. S1A). Only units recorded from shanks which passed through eYFP-labeled L6CT axons from V1 in the lateral, calretinin-negative zone of the pulvinar (23) were included for pulvinar analyses (e.g., second shank in Fig. 1C), while units recorded more medially were treated separately (e.g., first shank in Fig. 1C, Figs. S1B-D). While mice viewed either square-wave drifting gratings (“Visual trials”, see Methods) or a gray screen (“Blank trials”), ChR2-expressing cell bodies in V1 were stimulated with one second of sustained blue LED light at three different intensities (“low”, “medium”, and “high”, Fig. 1A; see Methods). This allowed us to further probe the possibility that different degrees of L6CT stimulation might account for the variety of results previously observed with sustained light delivery (3, 4, 15, 16). We also conducted V1 recordings in a subset of AAV-injected Ntsr1-Cre mice to verify that sustained light delivery was activating L6CTs (Figs. S2A-F).

Consistent with prior reports in anesthetized mice (3, 4), L6CT photostimulation with sustained light delivery at all intensities significantly suppressed visually-evoked firing rates (Figs. 1D-G; n=85 single-units from 7 shanks in 4 animals) in dLGN of awake mice (p<=0.003 for visual trials with vs. without LED in all light conditions, Wilcoxon signed-rank tests). Suppression of spontaneous firing rates was also significant with high-level light and approached significance in lower light level conditions (Figs. S1E-G; p=0.054, 0.056, 0.008 for blank trials with vs. without LED in low, med and high conditions). This effect is exemplified by the first example unit in Fig. 1D, whose visually-evoked and spontaneous firing rates were suppressed partially by low light and dramatically by high light. Nevertheless, there was heterogeneity in responses. For instance, some units (e.g., second example unit in Fig. 1D) were strongly activated by higher light levels and tended to be close to each other along the dorsal-ventral axis of dLGN (Fig. 1F), suggesting that the direction of their L6CT modulation may be related to their spatial (retinotopic) position within the nucleus. Yet overall, the majority of units in dLGN (65.88%) were significantly suppressed by L6CT photostimulation (Fig. 1G), suggesting that the dominant effect of sustained L6CT photostimulation on first-order visual thalamus dLGN is suppression.

In the pulvinar, we observed strikingly similar effects of L6CT photostimulation to what we observed in dLGN. The lateral pulvinar population (n=173 single-units from 10 shanks in 6 animals) was strongly suppressed by L6CT photostimulation during visual trials (Figs. 1H-K) as well as blank trials (Figs. S1H-J) at all light levels (p<0.001 for visual trials and blank trials with vs. without LED in all conditions; see first example unit in Fig. 1H), and the majority of lateral pulvinar units (53.76%) were significantly suppressed (Figs. 1K). In contrast, units recorded in the calretinin-expressing, medial area of pulvinar that lacks direct L6CT input from V1 (23) were considerably less visually-responsive or modulated by LED stimulation (Figs. S1A-D). As in dLGN, different light levels also had heterogenous effects in lateral pulvinar. For instance, the second example unit in Fig. 1H was activated by low light and suppressed by higher light levels. Interestingly, different light levels induced different changes in bursting in both dLGN and lateral pulvinar (Figs. S3A and S3C): significantly decreased bursting with low light (p<0.001 in dLGN and pulvinar, Wilcoxen signed-rank tests) and with medium light in dLGN (dLGN: p=0.002, pulvinar: p=0.098) and increased bursting with high light in lateral pulvinar but not in dLGN (dLGN: p=0.278, pulvinar: p<0.001). Light-induced changes in activity were not observed in dLGN or pulvinar units recorded from uninjected control animals (Figs. S1K-N), demonstrating that they were due solely to specific manipulation of L6CT activity. Thus, we demonstrate for the first time *in vivo* that visually-evoked and spontaneous activity in pulvinar, much like in anesthetized (3, 4) and awake dLGN (above), is suppressed by sustained L6CT photostimulation.

### Sustained L6CT photostimulation activates TRN

How do glutamatergic L6CTs exert inhibitory influences over their thalamic targets? We noticed that many units exhibited transient facilitation at the onset of L6CT photostimulation before subsequent suppression (e.g., first example unit in Fig. 1H; average PSTHs of suppressed cells in dLGN and pulvinar in Figs. 1G and 1K), suggesting that feed-forward inhibition is likely involved. Since TRN receives axon collaterals from L6CTs in V1 (11) and in turn provides GABAergic input to both dLGN and pulvinar (25), it is a likely source of feed-forward inhibition. TRN has been shown to be activated by L6CTs in slice (3, 18, 26, 27) and by L6CTs in auditory cortex *in vivo* (15), but how it is engaged by L6CT photostimulation in V1 of an awake animal has not been tested.

We recorded single-unit activity in the visual sector of the TRN (visTRN) with the same L6CT stimulation conditions as for dLGN and pulvinar recordings (Fig. 2A). As one would expect if recordings were well-targeted to visTRN (Fig. 2B), more than half (59.38%) of units were significantly visually-responsive, and many (62.50%) exhibited fast-spiking profiles, in contrast to in dLGN (11.76%) and pulvinar (1.73%).

**Fig. 2.**
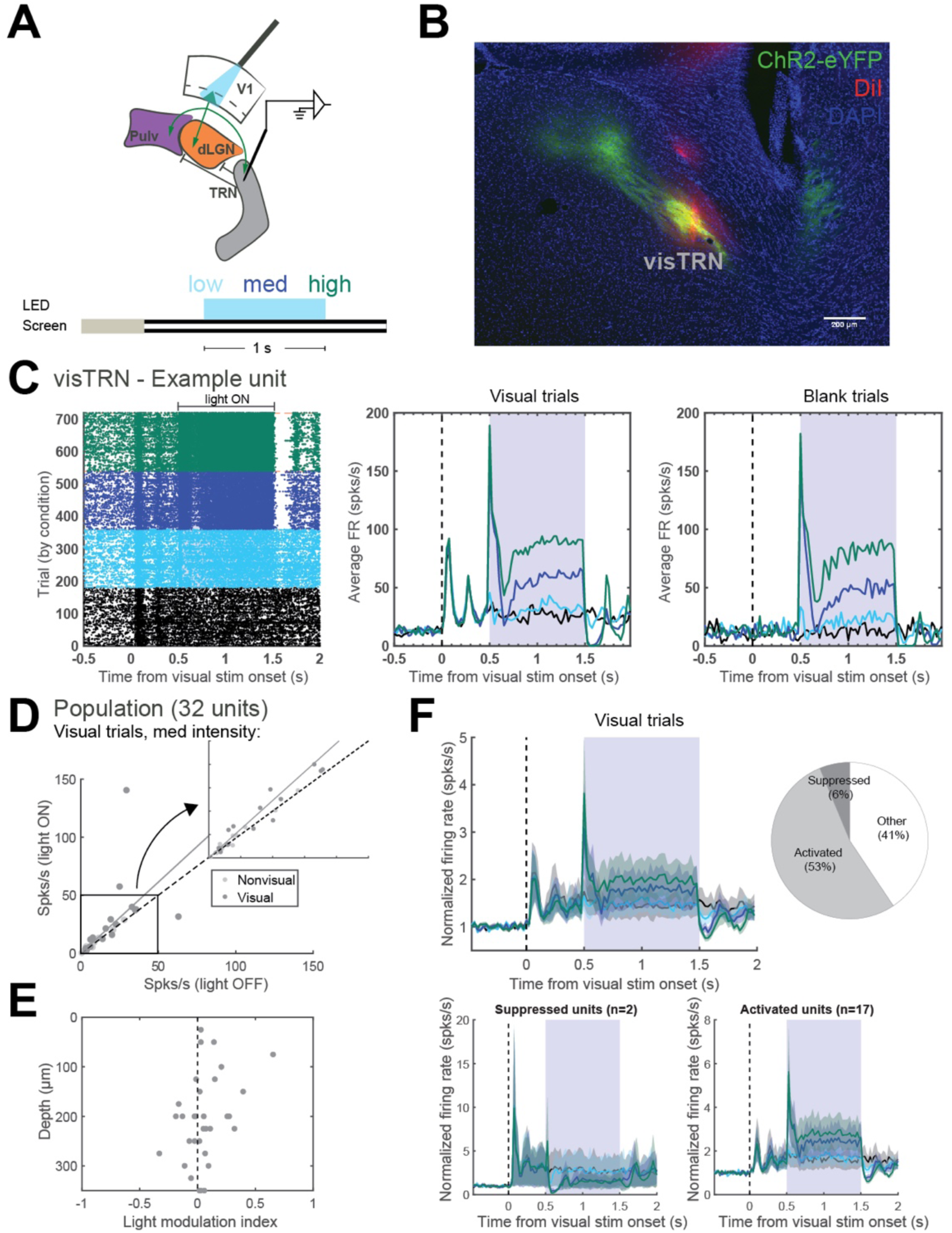
L6CT photostimulation with continuous light delivery activates units in visTRN. (A) Diagram of photostimulation and visTRN recording locations and trial structure for visual and LED stimulation. (B) Coronal section depicting ChR2-eYFP-positive axon terminals in the visual sector of the thalamic reticular nucleus (visTRN) from a Ntsr1-Cre mouse injected with AAV-DIO-ChR2-eYFP. The DiI track from the recording probe (red) overlaps with the L6CT axon terminal field. (C) Raster plot (left) and PSTHs from visual trials (middle) and from blank trials (right) for an example unit recorded in visTRN. (D) Average firing rates during the 1-second photostimulation period from visual trials, with versus without medium-intensity L6CT photostimulation. Inset: expanded scatter plot from area within the square (0-50 spks/s). (E) Light modulation index by depth within visTRN. (F) Effects of different photostimulation intensities on units’ visually-evoked activity in visTRN. Top, left: average normalized PSTH across all visTRN units. Top, right: proportion of units which were significantly suppressed or activated in two or more photostimulation conditions. Bottom: average normalized PSTH across suppressed (left) and activated (right) units. Shading indicates ±1 standard error of the mean.

Consistent with the hypothesis that dLGN and pulvinar suppression during sustained L6CT cell body stimulation is mediated by the visTRN, many units in visTRN were rapidly and strongly activated under these same conditions (Fig. 2C). At the population level (n=32 single-units from 4 penetrations in 3 animals), both visually-evoked and spontaneous firing rates were significantly increased with L6CT photostimulation at all light levels during visual trials (Fig. 2D; p=0.029, 0.023, 0.003 for low, med, and high LED) and approached significance in blank trials (p=0.014, 0.05, 0.054). Similar to observations in dLGN and pulvinar, the rate of bursting was also significantly reduced with low- and medium-intensity L6CT stimulation (Fig. S3C; p=0.021, 0.043, 0.34 for low, med, and high LED). Average visTRN population activity was increased by L6CT stimulation in a graded manner for the full duration of the photostimulation period, and the majority (53.13%) of units were significantly activated (Fig. 2F). Importantly, the activation latency of the TRN population by L6CT photostimulation (Fig. 2F) was the same as in dLGN (12ms; Fig. 1G), which argues that the TRN was engaged monosynaptically rather than indirectly by its reciprocal connections with dLGN and pulvinar. Therefore, since visTRN is rapidly activated during sustained L6CT photostimulation and provides GABAergic input to dLGN and pulvinar, it should provide disynaptic inhibition to dLGN and pulvinar under these conditions. This is consistent with our prior observations of suppressed firing rates in both nuclei (Fig. 1).

### L6CT axon terminal stimulation does not suppress dLGN and pulvinar

To further explore the necessity of TRN engagement for thalamic suppression during continuous L6CT photostimulation, we recorded from dLGN and/or pulvinar using a two-shank “optrode” (Figs. 3A and 3F; Methods) that delivers blue light at the same site as the recording contacts (21). We hypothesized that by directly photostimulating ChR2-expressing L6CT axon terminals within the dLGN and/or pulvinar instead of their cortical cell bodies, the TRN would not be directly engaged and thus inhibition of dLGN and pulvinar would be greatly reduced. Indeed, sustained (1-second) axon terminal stimulation elicited responses in dLGN and pulvinar that were very different from what we previously observed with cell body stimulation (Figs. 3B and 3G). In experiments that included at least one shank in dLGN (n=88 single-units from 5 shanks in 4 animals), light levels had to be carefully titrated because aberrant activity followed by extended periods of silence were observed if the light intensity was too high (Methods). This likely reflects the much higher density of L6CT axon terminals in dLGN versus pulvinar (12). Because of this, lower light levels were used in recordings that included dLGN, and in these instances the ramp-like increase in activity in pulvinar was largely absent (Figs. S4A-D). Under these stimulation conditions, effects on dLGN activity were variable (Fig. 3C; p=0.316, 0.003 and 0.498, visual trials with vs. without low, ramp to high and high LED, respectively; p=0.002, <0.001, 0.752, blank trials). In pulvinar-only experiments (n=129 single-units from 6 shanks in 3 animals), higher light levels could be used and resulted in enhancement of pulvinar activity (p=0.001, 0.067, <0.001, visual trials with vs. without low, med, and high LED; p=0.185, <0.001, <0.001, blank trials; Fig. 3H). As with the subset of facilitated units in cell body stimulation experiments in dLGN, units that were activated by L6CT axon terminal stimulation in both nuclei were also spatially clustered along the dorsal-ventral axis (Figs. 3D and 3I). These effects were not seen in an optrode recording from an uninjected control mouse (Figs. S4E-H), demonstrating that they were not due to the light itself or to damage from the optrode.

**Fig. 3.**
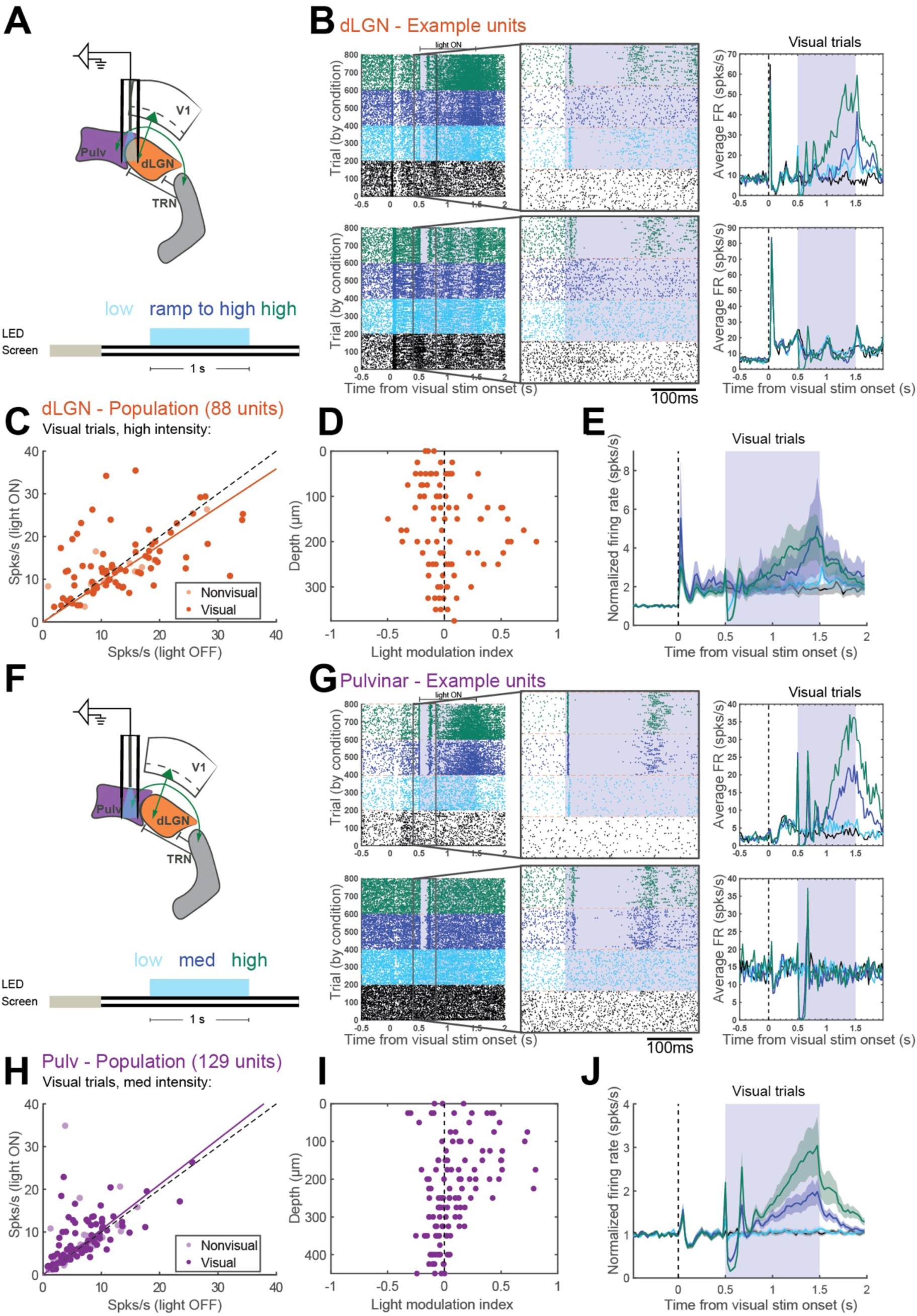
Photostimulation of L6CT axon terminals in dLGN/pulvinar does not suppress activity in visual thalamic nuclei. (A) Diagram of optrode configuration and trial structure for visual and LED stimulation. (B) Two example dLGN units. Left panels: raster plots. Middle panels: zoomed-in images of boxed parts of raster plots (from 100ms before to 300ms after LED stimulation onset). Right panels: PSTHs of average firing rates during visual trials. (C) Average firing rates during the 1-second photostimulation period from visual trials, with versus without high-intensity L6CT photostimulation. (D) Light modulation index by depth within dLGN. (E) Average normalized PSTH from visual trials across all dLGN units. Shading indicates ±1 standard error of the mean. (F-J) Same as (A-E) but for units recorded in lateral pulvinar, during experiments in which the optrode was entirely in the pulvinar and higher light levels were used (see Methods).

Overall, changes to visually-evoked and spontaneous population activity with L6CT terminal stimulation were similar between dLGN and pulvinar (Figs. 3E and 3J) and qualitatively very different from those with cell body stimulation (Fig. 1). Although some antidromic activation of cell bodies from axon terminal photostimulation may have occurred (28), the fact that we see such different effects on thalamic activity from when we photostimulated L6CT cell bodies strongly suggests that antidromic activation, if present, was weak. Instead, given that visTRN is activated by L6CT cell body photostimulation (Fig. 2) and direct L6CT axon terminal photostimulation in dLGN/pulvinar does not suppress (and can even excite) dLGN and pulvinar activity (Fig. 3), the potent net-suppression observed with sustained L6CT cell body stimulation is likely mediated by disynaptic inhibition through TRN.

### Frequency-dependent effects of L6CT photostimulation on dLGN and pulvinar

While our experiments with continuous light delivery for L6CT activation demonstrate L6CTs’ ability to suppress their first- and higher-order thalamic targets, can they also modulate thalamic activity in other ways? Continuous light delivery is a relatively uncontrolled method for photostimulation, and our own V1 recordings demonstrate variable effects of continuous light on L6 units’ activity that sometimes exceeded physiologically-relevant levels (Figs. S2C-F). To overcome some of the shortcomings of continuous LED stimulation and to test for possible frequency-dependent influences that have been described *in vitro* (18), we used a train stimulation paradigm (Fig. 4A; 10ms LED pulses at 1, 10, and 20 or 40 Hz for 1 second) to stimulate L6CT cell bodies during the same dLGN and pulvinar single-unit recording sessions from Fig. 1. We hypothesized that subsequent stimulation pulses in a 10Hz stimulation train, and perhaps also at higher frequencies, would produce increasing spike outputs as demonstrated in the somatosensory system *in vitro* (18).

**Fig. 4.**
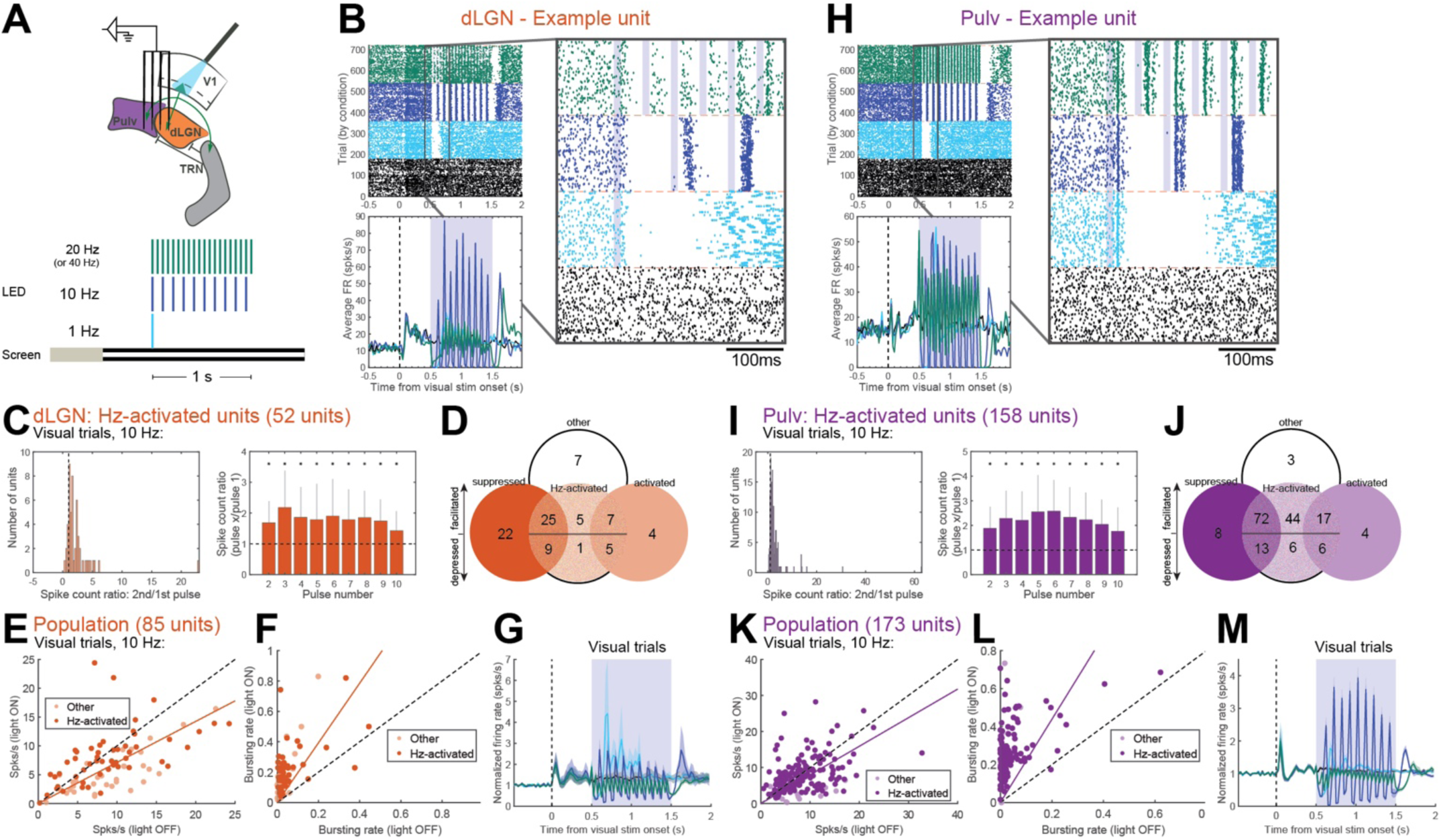
10Hz photostimulation of L6CTs facilitates spiking in dLGN and pulvinar. (A) Diagram of photostimulation and dLGN/pulvinar recording configuration and trial structure for visual and LED stimulation. (B) An example unit recorded in dLGN. Left: raster plot (top) and PSTH of average firing rates across visual trials (bottom). Right: zoomed-in image of boxed part of raster plot (100ms before to 300ms after LED photostimulation onset). Shaded rectangles indicate 10ms photostimulation pulses. (C) Quantification of facilitating spiking during 10Hz photostimulation trials across Hz-activated units in dLGN (see Methods for “Hz-activated” unit classification). Left: histogram of spike count ratios (spike outputs following the second photostimulation pulse relative to the first in a 10Hz train). Right: median spike count ratios across Hz-activated units, comparing spike outputs following photostimulation pulses 2-10 relative to the first pulse. Asterisks indicate ratios significantly different from 1 (p’s <0.05, sign test), and error bars indicate interquartile range. (D) Venn diagram of “suppressed”, “activated”, and “other” units (from 1s sustained photostimulation experiments, Fig. 1) whose spiking was facilitated (spike count ratio >1), depressed (spike count ratio <1), or not modulated (not “Hz-activated”) by 10Hz L6CT photostimulation. (E) Average firing rates during the 1-second photostimulation period from visual trials, with versus without 10Hz photostimulation. Saturated points indicate Hz-activated units included in quantification in (C). (F) Average bursting rates (number of spikes that occurred during bursts / total number of spikes) for all units during visual trials with and without L6CT photostimulation. (G) Average normalized PSTH from visual trials across all dLGN units. Shading indicates ±1 standard error of the mean. (H-M) Same as (B-G) but for pulvinar. Example unit in (H) is the same example unit as in Fig. 1H (top).

Indeed, in both dLGN and pulvinar, we consistently observed facilitating spiking following subsequent pulses in a 10Hz train (example units in Figs. 4B and 4H). Whereas a single photostimulation pulse elicited at most a weak and short-lived response, this response increased dramatically with further 10Hz stimulation pulses. The facilitation effect for each unit can be quantified by comparing the number of spikes following any pulse in the train to the number of spikes after the first pulse; thus, spike count ratios greater than 1 indicate facilitating spiking. Of the units recorded in dLGN and pulvinar that exhibited spiking responses to individual 10Hz light pulses (52/85 and 158/173 units considered “Hz-activated” in dLGN and pulvinar, respectively; Methods), the majority had spike count ratios (pulse 2/pulse 1) much greater than 1, and median spike count ratios were greater than 1 for all subsequent pulses (Figs. 4C and 4I; asterisks indicate p<0.01, sign test). These same signatures of facilitating spiking were absent from laminar recordings in V1 (Figs. S2J and S2L), indicating that this phenomenon is particular to L6 corticothalamic (as opposed to intracortical) synapses. Notably, the example pulvinar unit (Fig. 4H) is the same unit depicted in Fig. 1H (top); while it was strongly suppressed by sustained photostimulation, it exhibited facilitating spiking when driven by 10Hz photostimulation. In fact, many units in both thalamic nuclei that were significantly suppressed by sustained L6CT photostimulation exhibited spike facilitation (spike count ratio >1) from 10Hz photostimulation (25/56 and 72/93 in dLGN and pulvinar, respectively; Figs. 4D and 4J). While some thalamic units (like those depicted in Figs. 4B and 4H) also exhibited facilitating spiking at 20Hz, this was less consistent across units (Figs. S5A-C). This could be related at least in part to limitations imposed by ChR2 kinetics (29), though reductions in L6CT spiking were weak or absent under our experimental conditions (Figs. S2G-K).

The effects of moderate-to-high-frequency L6CT photostimulation on average firing rates across the full photostimulation period are, if anything, significantly suppressive in dLGN (Fig. 4E: p=0.07, <0.001, <0.001 for 1Hz, 10Hz and 20-40Hz in visual trials; p’s<=0.004 in blank trials) and approaching significance in pulvinar (Fig. 4K: p=0.001, 0.054, <0.001 in visual trials, p<0.001, 0.068 and <0.001 in blank trials) due to the temporally restricted and bursting nature of their spike outputs. In fact, the incidence of burst (as opposed to tonic) spikes increased dramatically under each train photostimulation condition in both dLGN (Figs. 4F and S3D, p’s <0.001 for 1Hz, 10Hz and 20Hz) and pulvinar (Figs. 4L, S3B and S3D; p’s <0.001). Therefore, moderate-frequency (10Hz) L6CT photostimulation profoundly alters the firing mode and facilitates the spiking responses of their thalamic targets (Figs. 4G and 4M) – even among those which are suppressed by the same pathway under different conditions.

Moderate-to-high-frequency train stimulation of L6CT axon terminals also had facilitating effects in dLGN (Figs. S5E-G) but not in pulvinar (Figs. S5H-J). However, facilitating effects may be even more difficult to identify with axon terminal stimulation, as ChR2 has been shown to incompletely recruit axons under moderate-to-high (10-40Hz) stimulation conditions and thus can lead to faulty indications of depressing synapses (30). Therefore, cell body stimulation experiments are most relevant to the question at hand, and they illustrate strongly facilitating spiking activity in dLGN and pulvinar.

### TRN engagement by L6CT photostimulation is also frequency-dependent

Prior *in vitro* work has shown that L6CT synapses onto TRN cells are also facilitating, yet the cumulative effect of the L6CT-TRN-thalamus pathway is depressing due to TRN’s intrinsic burst properties and depressing GABAergic inputs onto thalamic relay cells (18). Consistent with this, we observed facilitating spike outputs in visTRN recordings in response to 10Hz trains (Figs. 5A-C; p<0.001 at x=1:8, p<0.05 at x=9, spike count ratios pulse x/pulse 1). As with dLGN and pulvinar, 20Hz was not as effective at facilitating visTRN spiking (Fig. S5D), yet all conditions (1-20Hz) strongly increased the rate of bursting (Figs. 5F and S3D; p=0.001, p<0.001 and p=0.021 for 1Hz, 10Hz, and 20Hz).

**Fig. 5.**
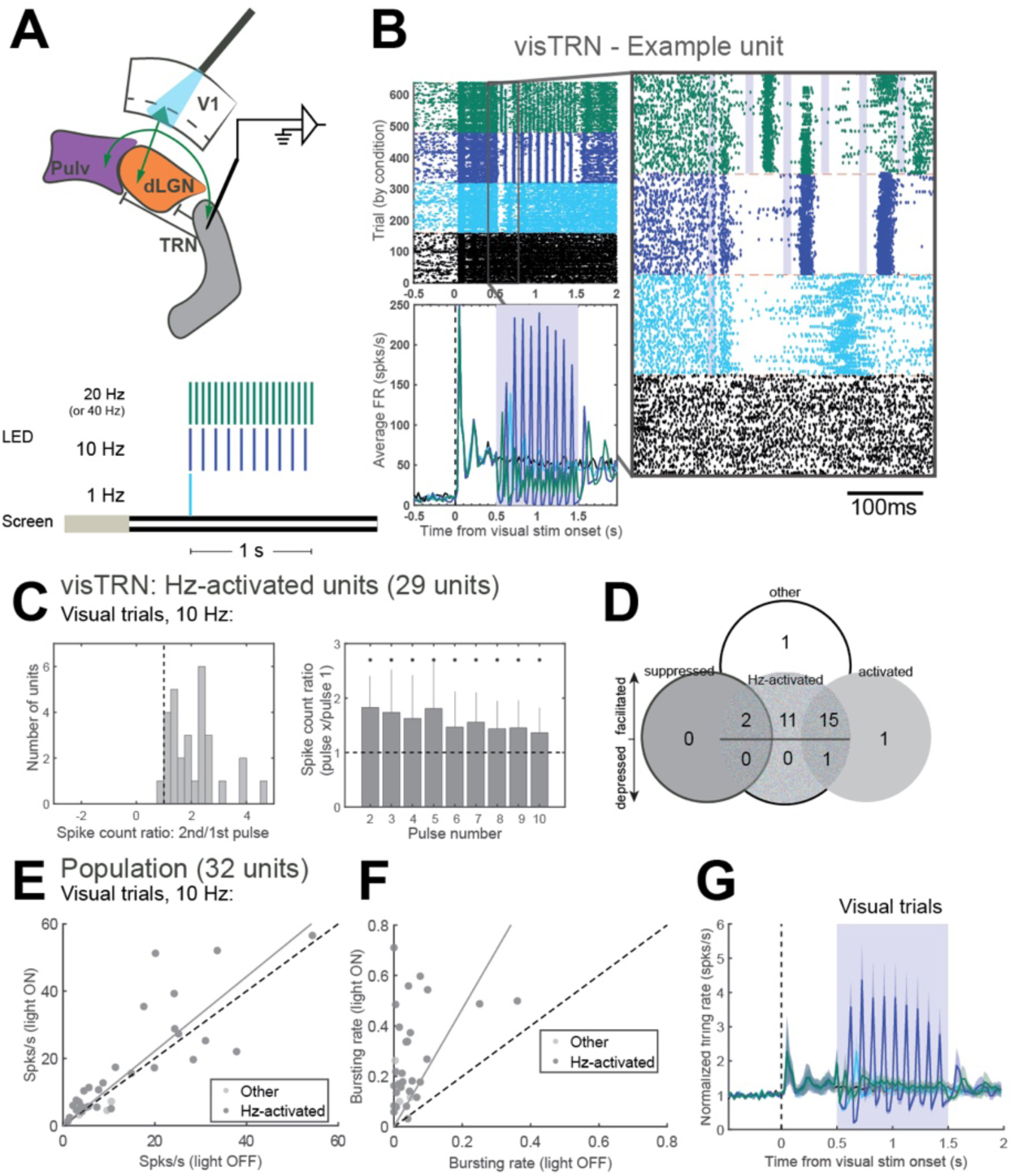
10Hz photostimulation of L6CTs also facilitates spiking in visTRN. (A) Diagram of photostimulation and visTRN recording configuration and trial structure for visual and LED stimulation. (B) An example unit recorded in visTRN. Left: raster plot (top) and PSTH of average firing rates across visual trials (bottom). Right: zoomed-in image of boxed part of raster plot (100ms before to 300ms after LED photostimulation onset). Shaded rectangles indicate 10ms photostimulation pulses. (C) Quantification of facilitating spiking during 10Hz photostimulation trials across Hz-activated units in visTRN. Left: histogram of spike count ratios (spike outputs following the second photostimulation pulse relative to the first in a 10Hz train). Right: median spike count ratios across Hz-activated units, comparing spike outputs following photostimulation pulses 2-10 relative to the first pulse. Asterisks indicate ratios significantly different from 1 (p’s <0.05, sign test), and error bars indicate interquartile range. (D) Venn diagram of “suppressed”, “activated”, and “other” units (from 1s sustained photostimulation experiments, Fig. 2) whose spiking was facilitated (spike count ratio >1), depressed (spike count ratio <1), or not modulated (not “Hz-activated”) by 10Hz L6CT photostimulation. (E) Average firing rates during the 1-second photostimulation period from visual trials, with versus without 10Hz photostimulation. Saturated points indicate Hz-activated units included in quantification in (C). (F) Average bursting rates for all units during visual trials with and without L6CT photostimulation. (G) Average normalized PSTH from visual trials across all visTRN units. Shading indicates ±1 standard error of the mean.

Overall, the cumulative effect of L6CT train stimulation on visTRN (Fig. 5G) is virtually identical to that on dLGN (Fig. 4G) and on the pulvinar (Fig. 4M). This is in contrast to the results of sustained cell body photostimulation experiments, where the cumulative effect on visTRN (Fig. 2F) was opposite of the majority of units in dLGN (Fig. 1G) and in the pulvinar (Fig. 1K). This similarity (rather than opposition) between dLGN/pulvinar and TRN suggests that with 10Hz train stimulation, activity in dLGN and pulvinar more closely reflects their direct, excitatory inputs while inhibition from the TRN is comparatively weak. In other words, these parallel monosynaptic excitatory and disynaptic inhibitory pathways from L6 to first- and higher-order thalamus are dynamically opposed, which can lead to highly flexible, context-dependent effects.

## Discussion

We set out to investigate how layer 6 corticothalamic neurons (L6CTs) influence first- and higher-order thalamus, using the visual system as a model. While these corticothalamic neurons are thought to serve modulatory roles in sharpening sensory responses and enhancing thalamocortical transmission in first-order thalamus (31), how they might accomplish such functions through their different excitatory and inhibitory routes to their thalamic targets is unclear. Moreover, the nature of the pathway from layer 6 to higher-order thalamic nuclei, like the mouse pulvinar, has not previously been explored, leaving many unanswered questions as to how these corticothalamic neurons might exert similar or dissimilar modulatory control over different classes of thalamic nuclei.

Using high-density multielectrode recordings with optogenetics in awake, head-fixed mice, we find very similar effects on both dLGN and pulvinar with L6CT photostimulation, but even within nuclei these effects vary drastically with the manner and degree of stimulation. Sustained optogenetic activation of L6CTs in V1 with different levels of continuous light strongly suppresses visually-evoked and spontaneous activity in dLGN and pulvinar, yet controlled 10Hz stimulation of this same population leads to facilitating spiking in both areas. These firing rate changes were accompanied by changes in burst versus tonic modes of firing, which is consistent with previous reports that corticothalamic feedback can modulate thalamic firing mode (6, 9, 10, 32). Remarkably, we also observed similar facilitating spiking at 10Hz in TRN, yielding virtually indistinguishable effects between TRN and pulvinar/dLGN. This stands in stark contrast to the effects on TRN and pulvinar/dLGN under sustained stimulation conditions, which were opposite in sign. These findings demonstrate the highly dynamic nature of these connections, whereby the relative balance between excitatory and inhibitory input to pulvinar and dLGN and the mode of thalamic firing can shift depending on the context of corticothalamic engagement. Therefore, describing the overall effect of the L6 corticothalamic pathway as simply suppressive or excitatory would fail to capture the functional nuance of this circuit.

### Effects of L6CT activation depend on the degree and manner of their activation

Previous optogenetic experiments using Ntsr1-Cre transgenic mice under anesthesia with continuous light delivery describe a net-suppressive influence of L6CT activation on first-order visual thalamus dLGN (3, 4). We replicate these findings in awake mice and show similar effects in the pulvinar. However, other effects of L6CT optogenetic stimulation have been reported in other species and other sensory systems; in particular, no change in dLGN firing rate in anesthetized ferret (8) and increased activity in first-order auditory and somatosensory thalamic nuclei MGBv and VPm, respectively (15, 16). While these discrepant findings could plausibly be attributed to differences between species and/or sensory systems, they could also be caused by methodological incongruities. Since the majority of these studies used uncontrolled, continuous light delivery for L6CT photostimulation, the intensity and duration of light stimulation, as well the level of viral expression, could all influence their reported observations. For instance, although we found significant population-level suppression of thalamic activity with each of our chosen light intensities, there was some heterogeneity such that some units’ activity changed in opposite directions with different light intensities (e.g., bottom examples unit in Fig. 1D and 1H). Thus, in combination with our train stimulation experiments with the same population of neurons, our results can help reconcile some of these prior conflicting findings by illustrating how the complex nature of L6 corticothalamic circuitry can lead to very different downstream effects under different contexts of L6CT activity.

Another important factor that may determine how L6CTs influence their thalamic targets is their topographic alignment. A limitation of ours and other rodent L6CT optogenetic studies (3, 4, 10, 15, 16) is the use of broad ChR2 expression and photoactivation of many L6CTs across retinotopic, tonotopic, or somatotopic locations. Some evidence suggests that inhibition is broader than topographically-aligned excitation in cat dLGN (33) and rat VPm (34). Thus, it is entirely possible that our broad activation of L6CTs throughout V1, our use of full-field visual stimuli and simultaneous single-unit recordings from different retinotopic locations in thalamus may have biased us toward observing suppressive effects under sustained L6CT photostimulation conditions. Future studies, perhaps in other species (e.g., ferret, non-human primate) with larger cortices and finer retinotopic maps or with more sophisticated methods for targeted optogenetic stimulation, could directly test the possibility that the relative balance between monosynaptic excitatory and disynaptic inhibitory pathways *in vivo* also depends on the retinotopic alignment between corticothalamic partners.

### Potential mechanisms underlying the dynamic nature of corticothalamic pathways

A major strength of this study is our investigation of how L6CTs impact activity in their first- and higher-order thalamic targets under a variety of different conditions in an awake animal. Our combination of recordings in dLGN, pulvinar and visTRN have allowed us to explore how these circuit components are recruited under various photostimulation conditions, yet we are limited by our *in vivo* extracellular recording methodology in capturing all possible mechanisms underlying our observed effects. For instance, the patterns of activity we see are also likely to be influenced by the reciprocal excitatory/inhibitory connections between the TRN and both dLGN and pulvinar (35, 36). Thalamic inhibition could also come from local inhibitory interneurons which, though quite rare in the rodent thalamus and especially in the pulvinar (37), are synaptically connected to L6CTs (26, 38). Nevertheless, the lack of pronounced suppression with axon terminal (Fig. 3) compared to cell body photostimulation (Fig. 1) suggests that they are not sufficient for the suppressive effects we observe under the latter condition.

Meanwhile, the consistency of our findings with prior *in vitro* work (18) allow us to speculate on likely synaptic mechanisms underlying the frequency-dependent effects we observe. First, short-term facilitation in L6CT synapses is a robust and widely described phenomenon and is an important component of frequency-dependent spiking described in the somatosensory slice preparation (18). Excitatory synaptic inputs from L6CTs to dLGN neurons have also been shown to persistently facilitate with moderate-to-high frequency (up to ∼25Hz) electrical stimulation of the corticogeniculate pathway *in vitro* (39, 40) and in the anesthetized cat (41), as well as to pulvinar neurons *in vitro* (13). Based on our observations of spike facilitation in the TRN with 10Hz L6CT photostimulation, which is consistent with the known facilitating nature of these synapses (3, 18, 26, 27), one might have predicted that facilitating inhibition from TRN would balance with facilitating excitation from L6CTs to produce consistent (rather than facilitating) spike outputs in dLGN and pulvinar. Instead, we also observed robust spike facilitation in dLGN and pulvinar. One possible explanation is that L6CT-TRN facilitation is weaker than at L6CT-relay cell synapses (18, 26, 27), which would lead to net-facilitation of the monosynaptic excitatory pathway overall. Another potential explanation is that GABAergic synapses from TRN to thalamic relay neurons exhibit prominent synaptic depression (18), which would allow the relative influence of disynaptic inhibition through TRN to weaken overall with higher-frequency L6CT activity. Thus, we suspect that facilitating excitation and depressing inhibition through TRN are both important synaptic mechanisms underlying our frequency-dependent effects.

Still other mechanisms could also be at play that are not mutually exclusive with the previously described circuit and synaptic contributions. For instance, thalamic neurons’ intrinsic properties, and in particular the presence of T-type calcium channels which lead to bursting when “deinactivated” at hyperpolarized membrane potentials (42, 43), may also contribute to the frequency-dependent effects we see. In fact, we observed a pronounced shift towards increased bursting during train photostimulation experiments (Figs. 4F, 4L, 5F, and S3D). These intrinsic properties likely account for the characteristic rebound spiking (and increased bursting) we observed in all three thalamic nuclei following ∼100ms of silenced activity with 1Hz photostimulation (Figs. 4B, 4H and 5B), which corresponds to the approximate time for T-type calcium channels to become deinactivated by hyperpolarizing input (from TRN, for instance) and trigger a shift into burst mode (42, 43). Still, we hypothesize that these intrinsic properties are not solely responsible for the spike facilitation we observe with 10Hz photostimulation because higher-frequency (20Hz) photostimulation also increased bursting (Fig. S3D) but did not produce spike facilitation (Figs. S5B-C). Moreover, thalamic relay neurons *in vitro* have been shown to exhibit spike facilitation (primarily within the first few L6CT stimulation pulses, as was the case in our experiments) even when held at hyperpolarized membrane potentials to induce a sustained burst mode of firing (18). To what extent these intrinsic properties contribute to our frequency-dependent effects could be tested directly in mice with T-type calcium channels genetically deleted (44). Overall, we suggest that multiple mechanisms (intrinsic physiological, synaptic, and circuit) all contribute to the highly dynamic nature of L6CT corticothalamic pathways.

### L6CT influences on first- versus higher-order thalamus

This study offers the first investigation of how L6CTs influence their higher-order thalamic targets, such as the pulvinar in the rodent visual system, *in vivo*. We find that, overall, L6CT photostimulation had similar effects on lateral pulvinar as on first-order thalamus dLGN that were also highly dynamic and frequency-dependent. These findings are consistent with hypothesized functional similarities between L6CT projections to both classes of thalamic nuclei based on their similar “modulator”-like morphological and physiological characteristics (1, 45). Nevertheless, there are some differences between L6CT projections to dLGN and pulvinar that could have led to functional disparities. For instance, while many L6CTs that project to dLGN also project to the pulvinar (11), those that project to both classes of thalamic nuclei are only found in lower L6 (11, 46–48). The pulvinar receives additional L6 input from other visual cortical areas, and pulvinar-projecting L6CTs in these areas are found throughout L6 (46). Thus, by restricting photostimulation to V1, we are engaging a subset and perhaps even a minority of the pulvinar’s L6CT inputs. Moreover, a recent study identified a small population of L6B cells that project exclusively to higher-order thalamus, but not to first-order thalamus or TRN (49); if and how this population’s functional influence on pulvinar differs from that of the more common L6CT population would be of significant interest. Yet despite these anatomical differences, we see remarkably similar effects of L6CT photostimulation in the pulvinar and dLGN that may be indicative of fundamentally similar functions for these corticothalamic pathways.

Another distinguishing feature of higher-order compared to first-order thalamic nuclei is that L6CTs are not their only source of corticothalamic input. Higher-order nuclei like the pulvinar also receive CT projections from layer 5 (L5), and these are hypothesized to act as the primary “driving” inputs in lieu of strong input from the sensory periphery (1, 45). Given the suppressive influence of L6CTs on other cortical populations within V1 (3), it is therefore possible that the inhibitory effects we see in pulvinar with sustained L6CT photostimulation are due not only to the engagement of the visTRN as we describe, but also to indirect suppression of the L5CT “driving” inputs. While we cannot rule out this possibility, we note that L5, including lower L5 (“L5B”) where subcortically-projecting L5 neurons in V1 are somewhat biased to reside (50), was not fully inactivated under our light stimulation conditions (Fig. S2D). Moreover, the fact that we see such similar and robustly facilitating spiking with moderate-frequency L6CT photostimulation in pulvinar as in dLGN cannot easily be explained by a mechanism involving L5CTs. Instead, we believe our findings provide compelling *in vivo* evidence of functional similarity in L6CT projections to different classes of thalamic nuclei.

### Summary

Our results contribute to a broader understanding of the circuit computations underlying L6CTs’ functions in both first- and higher-order sensory thalamus.Observations of visual response suppression by L6CTs in dLGN (3, 4) has led some to suppose that this is the primary mechanism by which these corticothalamic neurons wield functional influence over their thalamic targets. However, our results suggest that L6CTs take a more nuanced approach. We propose that L6CTs can exert net-inhibition over their thalamic targets under some conditions – such as when only transiently or weakly activated – and in others can strongly facilitate their targets’ responses. Further, these connections can regulate not only the level but also the mode of thalamic activity, as demonstrated by our observed changes in burst versus tonic firing under different photostimulation conditions. Our data suggest that this functional flexibility is mediated by careful balancing between the parallel excitatory and inhibitory (through TRN) routes from cortical layer 6 to dorsal thalamus, altogether allowing L6CT projections to flexibly control thalamocortical input. Since we see similar L6CT effects in both first- and higher-order visual thalamus, we expect these projections to affect not only thalamocortical transmission to V1, but also cortico-thalamo-cortical transfer through the pulvinar to higher-order visual cortical areas. These dynamic corticothalamic pathways could afford numerous functional and computational advantages, such as for stimulus-specific amplification (31, 39) and even higher-level representations, like perceptual decision confidence in the pulvinar as demonstrated by computational modeling of these pathways (51). With an improved understanding of the highly dynamic nature of these L6CT pathways to first- and higher-order thalamus in an awake animal, future work may be able to elucidate their role in modulating sensory processing in the context of sensory-guided behaviors

## Materials and Methods

### Animals

Male and female Ntsr1-Cre GN220 transgenic mice (GENSAT) aged 8-14 weeks (with the exception of three mice, which were 16-17 weeks old) were used for experiments. Cre-negative animals were used for control experiments. All experimental procedures followed protocols approved by the Salk Institute Animal Care and Use Committee.

### Surgeries

Mice were first anesthetized with a ketamine/xylazine cocktail (100mg/kg of ketamine and 10mg/kg xylazine) via intra-peritoneal injection and then placed in a stereotax (David Kopf Instruments Model 940 series). A small craniotomy was made over primary visual cortex of the left hemisphere (coordinates relative to bregma: 3.20mm posterior, 2.65mm lateral). A total of 100-150nl of AAV5-EF1a-DIO-hChR2(H134R)-eYFP was pressure-injected via Picospritzer (General Valve Corp) or syringe through a 25-30μm pipette at 1-2 depths, 0.3-0.6mm from pial surface. Viruses were injected at an approximate rate of 20nl/min, and the pipette was left in place for at least 5 minutes following injection prior to removal. Mice were returned to their cages and given 2.5-3 weeks before experimentation.

Four to seven days before experimentation, mice underwent an acute surgery for headframe implantation. Skin was cleared away so that a circular headframe (7mm inner diameter) could be attached with dental cement (C&B-Metabond, Parkell). A dull pipette attached to a micromanipulator (MP-285, Sutter Instrument Co) was used to relocate bregma and mark positions with a waterproof pen for targeting thalamus recordings (coordinates relative to bregma: 1.25-2.75mm lateral, 1.8-1.9mm posterior). The skull was covered with a silicone elastomer (Kwik-Cast, World Precision Instruments) and mice were given a carpofen subcutaneous injection (5mg/kg), Ibuprofen in their water bottles and at least 24 hours undisturbed in their cages.

### In vivo electrophysiology

Prior to recordings, mice were given 2-4 training sessions to habituate to the running wheel. One day prior and on the day of recordings, mice were given a dilution of dexmethasone (15mg/kg) to alleviate brain swelling. On the day of recording, mice were anesthetized with isoflurane, and a craniotomy was made over the thalamus of the left hemisphere. For cell body stimulation experiments, an additional craniotomy was made over the injection site in V1 (in two animals the cortex was heavily thinned). Mice were then head-fixed on a wheel, where they were free to run at their will and movement was tracked with a rotary encoder. Silicon microprobes (21) were coated with a 2.5-5% solution of DiI (D282, ThermoFisher) in distilled water or ethanol and lowered into thalamus with a micromanipulator (MP-285, Sutter Instrument Co). Probe configurations used for dLGN/pulvinar recordings were 128DN and 128D (128 channels across 4 shanks, 775μm vertical extent of electrodes, 150μm or 330μm separation between shanks, respectively) or 64G (64 channels across 2 shanks, 300μm separation between shanks, 525μm vertical extent of electrodes). A 64D probe (64 channels on one shank, 1.05mm vertical extent of electrodes) was used for visTRN recordings. For cell body stimulation experiments, a 1mm diameter optical fiber (1mm diameter, 0.39 NA, ThorLabs) was positioned at approximately a 50-60° angle from and 0.5-1mm above the surface of the V1 craniotomy. Multi-shank probes for thalamic recordings were oriented horizontally (medial-lateral). After the probe penetrated the cortical surface, agarose (2.5-3.5%; A9793, Sigma-Aldrich) was poured over to fill the well of the headframe holder, thus covering the probe shank(s) and the tip of the optical fiber. The probe was continuously lowered slowly down to ∼2.4-2.6mm beneath the cortical surface over the course of approximately 20 minutes. Once the probe was in its final position, it was allowed to sit and settle for 30 minutes before any data acquisition commenced. Data from all but two animals was acquired at 20kHz with an OpenEphys acquisition system (52), connected to an Intan RHD2000 128-channel amplifier board. Data from the remaining two animals was acquired at 20kHz with an Intan RHD2000 USB interface board.

### Visual and optogenetic stimulation

Visual stimuli was generated through custom MATLAB code using Psychtoolbox, as described previously (53) and presented on a 24” LED monitor (GL2450-B, BenQ). The monitor screen was positioned 12cm from the mouse’s right eye. Visual stimuli consisted of square-wave drifting gratings at four orientations in eight directions, 0.04 cycles/° spatial frequency, and 2Hz temporal frequency (one experiment in one mouse at 1Hz). A full “trial” consisted of a 0.5-second pre-stimulus period (grey screen), 2 seconds of visual stimulus presentation, and 1.5-2 seconds post-stimulus period (grey screen). 20% of trials were “blank” trials, in which the screen remained grey for the full trial duration.

Optogenetic stimulation was controlled via an Arduino Zero microcontroller board with a 12-bit DAC output pin. The Arduino interfaced with the visual stimulus generator code through a serial port connection to read in input parameters for start time, duration, intensity and frequency of LED stimulation from a custom Matlab GUI. All experiments consisted of a “no light” condition plus three different light conditions: sustained photostimulation at low, medium, or high light intensities, or trains of 10ms photostimulation pulses at 1, 10, and 20 Hz (40 Hz in two mice). All recorded units underwent both sustained and trains photostimulation experiments (except V1 units from two mice, which only underwent sustained photostimulation experiments). LED stimulation always lasted for 1 second and started 1 second after the trial start (0.5 seconds after visual stimulus onset). For cell body stimulation experiments, blue LED light was delivered through a custom optical fiber patch cord (1mm diameter, 0.39 NA, ThorLabs) connected to an LED driver (PlexBright LED 465nm, Plexon). For axon terminal stimulation experiments, blue light was delivered through a custom “optrode” (64G probe, integrated with a 200μm diameter, 0.22 NA multimode fiber halfway between the two shanks and ending just above the top contacts) connected to an LED driver (470nm fiber-coupled LED with T-Cube LED driver, ThorLabs). Light intensity was controlled by assigning output values for the Arduino DAC 12-bit output pin (0-4095). These output values were typically kept consistent across experiments, but power output was also measured with a power meter (PM100D with S121C power sensor, ThorLabs) as verification of consistency. Output power ranges from the Plexon LED through the 1mm diameter fiber were 0.7-1mW (“low”), 3.5-4.5mW (“med”), and 6.5-8.5mW (“high”). Output powers from ThorLabs LED to optrode with 200μm diameter fiber were estimated with a dummy fiber-optic implant (200μm diameter, 0.22NA). In these optrode experiments, DAC outputs assigned to “low”, “med” and “high” conditions had to be varied by experiment because it was observed in some experiments, and specifically in experiments in which at least one shank on the probe was in dLGN, too high light power caused a massive ramp-like increase in activity before units went completely silent for >20 seconds. Thus, light levels were titrated during recording so that this seemingly excitotoxic effect did not occur. Consequently, estimated light levels during experiments that included at least one shank in dLGN ranged from 120-152μW (“low”) and 250-560μW (“high”) as measured through the 200μm diameter dummy fiber. In these experiments, the third light condition was a 1 second ramp, in which the Arduino DAC output was linearly increased in bits over 1 second so that it ended at the “high” light level. In pulvinar-only optrode experiments, estimated light levels ranged from 120-152μW (“low”), 250-310μW (“med”) and 500-600μW (“high”).

### Histology

After recordings were completed, animals were given an intraperitoneal injection of euthasol dilution (15.6mg/ml) and then perfused with phosphate-buffered saline (PBS) followed by 4% paraformaldehyde. Brains were dissected out from skulls and post-fixed in 2% PFA and 15% sucrose solution at 4°C for ∼24 hours before being moved to 30% sucrose at 4°C for at least another ∼24 hours. Brains were frozen in sucrose and sliced on a freezing microtome into 50μm sections, starting from the anterior edge of the hippocampus to the posterior end of cortex. All sections were counterstained with 10μM DAPI in PBS for 10 minutes before being mounted and coverslipped with Polyvinyl alcohol mounting medium containing DABCO. Additional immunohistochemistry was performed on thalamic sections by incubating at 4°C for 16-20 hours with rabbit anti-calretinin primary antibody (1:1000; Swant 7697) in 1% Donkey Serum/.1% Triton-X 100/PBS, followed by donkey anti-rabbit conjugated to Alexa 647 (1:500; A-31573, Life Technologies) before DAPI counterstaining. Imaging was performed with an Olympus BX63 microscope with a 10X objective.

### Spike sorting

Spike-sorting on extracellularly recorded data was performed semi-automatically using Kilosort (54). Briefly, different recordings from the same recording session (e.g., different intensities experiment followed by trains stimulation experiment) were concatenated together into a single binary file, common average referenced, bandpass filtered (300-2000Hz) and spatially whitened prior to template-matching. Because onset and offset of LED stimulation for optogenetics caused artifacts that could be mistaken for spikes by the algorithm, spikes within 3ms of the light onset and offset were removed, and additional “spikes” were removed whose amplitudes surpassed 20 times the standard deviation of spike amplitudes. Clusters were further curated manually using Phy (55). Additional optogenetic artifacts were evident as points far outside the cluster of principal component features whose cross-correlograms with their assigned clusters exhibited clear refractory period violations and thus could be manually removed. For a few experiments in which the timing of the ADC input to OpenEphys from the Arduino (for controlling the LED) was not saved properly due to a bug in the OpenEphys GUI, thus prohibiting us from removing optogenetic artifacts based on their timing relative to LED onset and offset, we removed artifacts based only on their large amplitudes in Kilosort and as outliers in their principal component features in Phy. If artifacts were not removed, they were extremely obvious in unit raster plots because they lined up precisely with LED photostimulation onset/offset and thus were readily distinguishable from legitimate spikes, even those with short latency. Thus, we are very confident that we successfully removed the vast majority of optogenetic artifacts, and all short-latency responses that we report come from legitimate spikes. We did our best to also remove optogenetic “hash” from ChR2-expressing axons in our thalamic recordings, which usually manifested as points well outside the main cluster in the principal component features view that exhibited substantial refractory period violations in their cross-correlograms. These “hash”-y units were especially prominent in dLGN and, when analyzed separately, their spikes occurred nearly exclusively during photostimulation periods, further suggesting that they were not true thalamic units (which had high spontaneous firing rates). Spike clusters were manually assigned to noise, multi-unit, and good/single-unit categories, and only good single-units that had fewer than 0.5% refractory period violations and “unit quality” (isolation distance; method from (56), MATLAB code adapted from https://github.com/cortex-lab/sortingQuality) greater than 16 were included for analyses.

### Assigning units to thalamic nuclei

Prior to *in vivo* recordings, recording probes were coated in DiI and were easily visualized post-mortem. Staining for calretinin (described above) provided a clear border between lateral pulvinar (calretinin-negative) and dLGN (calretinin-positive axons from retinal ganglion cells), thus allowing for definitive assignment of units recorded on different probe shanks to lateral pulvinar or dLGN. In rare instances in which a DiI trace fell ambiguously right on the border between pulvinar and dLGN, units recorded from the corresponding shank were excluded from analyses. Calretinin staining also provided an additional border between lateral pulvinar and rostral-medial and caudal-medial pulvinar, where cell bodies were calretinin-positive. Pulvinar units were assigned to lateral pulvinar if the DiI trace from their corresponding probe shank passed through the calretinin-negative portion of pulvinar and through eYFP-positive axon terminals, whereas shanks passing mainly through calretinin-positive cells and absent or sparse eYFP-expressing axons were considered in medial pulvinar. Similarly, data from one shank in one experiment which passed through dLGN but not through eYFP-expressing axons were excluded from dLGN analyses. Calretinin staining did not allow us to distinguish between rostromedial and caudomedial pulvinar (23).

To approximate where the dorsal/ventral boundaries of thalamic nuclei (dLGN, pulvinar, visTRN) fell on each recording shank, we determined the anatomical boundaries of visually responsive units. To classify visually responsive units, we identified the visual direction which maximally changed each unit’s firing rate from baseline and used the spike counts from those preferred visual stimulus trials without LED stimulation to assess whether each unit’s activity was significantly different from “blank” trials without LED stimulation (Wilcoxon rank-sum test). We separately compared spike counts between blank and preferred-visual trials during the first 100ms of the visual stimulus (to assess visual onset response) and 0.5-1.5s from the visual stimulus onset (to assess sustained visual stimulus response), and thus considered units visually-responsive if either of these p-values passed the 0.025 threshold (Bonferroni correction for two comparisons). We then estimated the dorsal/ventral boundaries of visual thalamic nuclei as the first and last channels on each probe shank which had visually responsive units. These boundaries were corroborated across experiments from the same recording sessions, and if there were discrepancies the boundaries were set as those channels which had significantly visually responsive units in both experiments. In this way, all units analyzed from sustained photostimulation experiments were also analyzed for the trains experiments and vice versa. For visTRN recordings, we sometimes recorded visually-responsive units very far ventral and with very short latency, possibly through the optic tract. Thus, we also included as criteria that channels with visually-responsive units that were separated by more than 100μm from the nearest visually-responsive channel were excluded, and for visTRN specifically that depth of included channels should not exceed 400μm. Units were considered fast-spiking (FS) whose waveform trough-to-peak times were less than 0.4ms but were not analyzed separately because they were very uncommon in dLGN and extremely rare in pulvinar, consistent with other reports (17).

### Quantification and statistical analysis

Average firing rates were calculated from the one-second period of photostimulation (0.5-1.5s from visual stimulus onset). Although our mice did not run very often, we exclusively used stationary trials (speed < 2cm/s) for our average firing rate calculations since running has been shown not only to affect firing rates in V1 (57), but also in dLGN (58). Firing rates were calculated separately for visual and blank trials and for each light condition (no light, low, med, high).

For population-level statistics on the effects of L6CT photostimulation on these average firing rates, we pooled together all units from the same experiment type (e.g., trains vs. sustained photostimulation) and from the same thalamic nucleus across animals and recording shanks and compared their firing rates in trials without L6CT photostimulation to firing rates in each photostimulation condition using the Wilcoxon signed-rank test. Population-averaged peristimulus time histograms (PSTHs) were constructed using individual units’ average firing rate across trials (visual and blank trials separately) in 25ms bins across the full trial duration (2.5s). We then normalized to each unit’s baseline firing rate (calculated from the 500ms prestimulus period) and averaged these normalized PSTHs across units. Response latencies were estimated from population-averaged PSTHs in 1ms bins from blank trials, and latency was taken as the first time point after the LED onset which surpassed 2 standard deviations above baseline.

Average firing rates were also used to compute a light modulation index for each unit. The light modulation index was computed as the difference between average firing rates in visual trials with and without L6CT photostimulation (separately for each photostimulation condition) divided by their sum. Thus, a light modulation index greater than 0 indicates increased activity with L6CT photostimulation while less than 0 decreased activity.

To determine the statistical significance of each unit’s response to L6CT photostimulation, we compared spiking responses during the one second photostimulation period on all trials with and without photostimulation (separately for each photostimulation condition) with a Wilcoxon rank-sum test. In order to summarize how individual units were affected by sustained L6CT photostimulation, units were considered “suppressed” if their activity was significantly suppressed (p<0.05 and light modulation index < 0) in at least 2/3 conditions, or “activated” if their activity was significantly increased (p<0.05 and light modulation index >0) in at least 2/3 conditions.

To analyze frequency-dependent effects, we first sought to identify units that spiked in response to individual photostimulation pulses. This is because it was expected that not every neuron we recorded from should be directly innervated by our ChR2-expressing L6CTs (as demonstrated by non-uniform expression of ChR2-eYFP across each thalamic nucleus). With each unit’s peristimulus time histogram of spike counts (in 5ms bins) for visual trials with 10Hz or 20Hz photostimulation, we used the following criteria for so-called “Hz-activated” units: 1) at least one bin surpasses threshold (three standard deviations above the mean visually-evoked response, taken from the one-second photostimulation period in no light trials) within 50ms of at least the first or second photostimulation pulse in the train; 2) if there was no significant response to either of the first two photostimulation pulses, there had to be significant response bins (above threshold) following at least half of all subsequent pulses. Only these “Hz-activated” units were analyzed for spike count ratios, which were calculated as the total number of spikes within 50ms of each photostimulation pulse divided by the total number of spikes within 50ms of the first pulse. Since the distributions of these data were clearly non-normally distributed, we chose to plot the medians and interquartile ranges. The sign test was used to test the null hypothesis that the median of spike count ratios was equal to one (i.e., no change in spike outputs).

To investigate the effects of our different L6CT photostimulation conditions on thalamic bursting, for each unit we analyzed the interspike intervals (ISIs) for all spikes that occurred within the 1-second photostimulation period (.5-1.5s post-visual stimulus onset) across visual trials. To control for possible changes in bursting induced by behavioral state rather than our L6CT manipulations (57), we restricted these analyses to trials in which the mouse was stationary as for our firing rate analyses (speed < 2cm/s). Spikes occurring in bursts were defined per convention (e.g., 59) as those which were preceded by an ISI ≥ 100ms and followed by an ISI ≤ 4ms (first spikes in a burst) and all subsequent spikes which were preceded by an ISI ≤ 4ms. The bursting rate for each unit was calculated as the ratio of the number of spikes occurring in bursts to the total number of spikes, calculated separately for each photostimulation condition. Changes in bursting under each photostimulation condition compared to the no-photostimulation condition were assessed statistically using the Wilcoxen signed-rank test.

## Data and software availability

Data and analysis code will be made available upon reasonable request.

## Acknowledgements

We thank Ashley Juavinett and Pamela Reinagel for comments on the manuscript, Sotiris Masmanidis and Kwang Lee for help with optrode assembly, and other members of the Callaway lab for helpful discussions. This work was supported by NIH grants F31 EY028853 (M.A.K.) and R01 EY022577 (E.M.C.).

## Contributions

Conceptualization, M.A.K. and E.M.C.; Methodology, M.A.K. (optogenetics and electrophysiology) and A.D.F. (histology); Investigation and Formal Analysis, M.A.K.; Data Curation (spike-sorting), M.A.K. and A.D.F.; Writing – Original Draft, M.A.K.; Writing – Review and Editing, M.A.K., A.D.F. and E.M.C.; Funding Acquisition, M.A.K. and E.M.C.; Resources and Supervision, E.M.C.

## SI Appendix

### Additional Methods

#### V1 recordings

In two mice, additional experiments were performed to record single-unit activity in primary visual cortex with a single (64D) recording probe, while a few mice (n=3) only underwent V1 recordings with a two-shank (128AN) recording probe. The same optogenetic and visual stimulation conditions were used as for thalamic recordings, but with an additional stimulus protocol consisting of alternating 2-second periods of screen-on and screen-off for 2 minutes. From these short recordings, we performed current source density (CSD) analysis by taking the second spatial derivative using CSDPlotter (1) of the low-pass-filtered (<1000Hz) local field potential during the transitions between screen-off and screen-on periods. For each column of each shank, we assigned channels to layer based on the following criteria: congruent channels which exhibited the fastest current sink (typically 2-3 channels per column, spanning 100-150μm) were assigned to layer 4; those that fell above this initial current sink as layer 2/3; those that exhibited an additional but delayed current sink ting approximately 150-300μm beneath the bottom of layer 4 as layer 5B; those in between layers 4 and 5B and layer 5A; and those below layer 5B as layer 6. Units were separated into fast-spiking (FS) and regular-spiking (RS) categories based on their waveform trough-to-peak times (<0.475ms considered FS) because this metric appeared to clearly separate all recorded units into two main clusters (Fig. S2B). Firing rates and light modulation indices were analyzed as for thalamic units.

To assess how our different photostimulation conditions affected the firing rates of putative L6CT cells, we took two different approaches (Figures S2E-F). First, we used the Stimulus-Associated spike Latency Test (2) to identify L6 RS units with statistically significant responses within 10ms of photostimulation onset. However, because of extensive interconnectivity among cortical pyramidal cells which can lead to short-latency spikes in cells synaptically connected to ChR2-expressing cells, we suspect that there are some false positives in this sample. Thus, we took an additional approach of simply identifying L6 RS cells which were positively light-modulated (light modulation index > 0 in at least two photostimulation conditions), based on the observation that L6CTs primarily inhibit other cortical pyramidal cells through their connections with fast-spiking interneurons (3). Because these two different approaches yielded very similar estimates of putative L6CT firing rates across our different photostimulation conditions, we felt that our phototagging approach using the SALT test provided a reasonable approximation of the L6CT population.

**Fig. S1.**
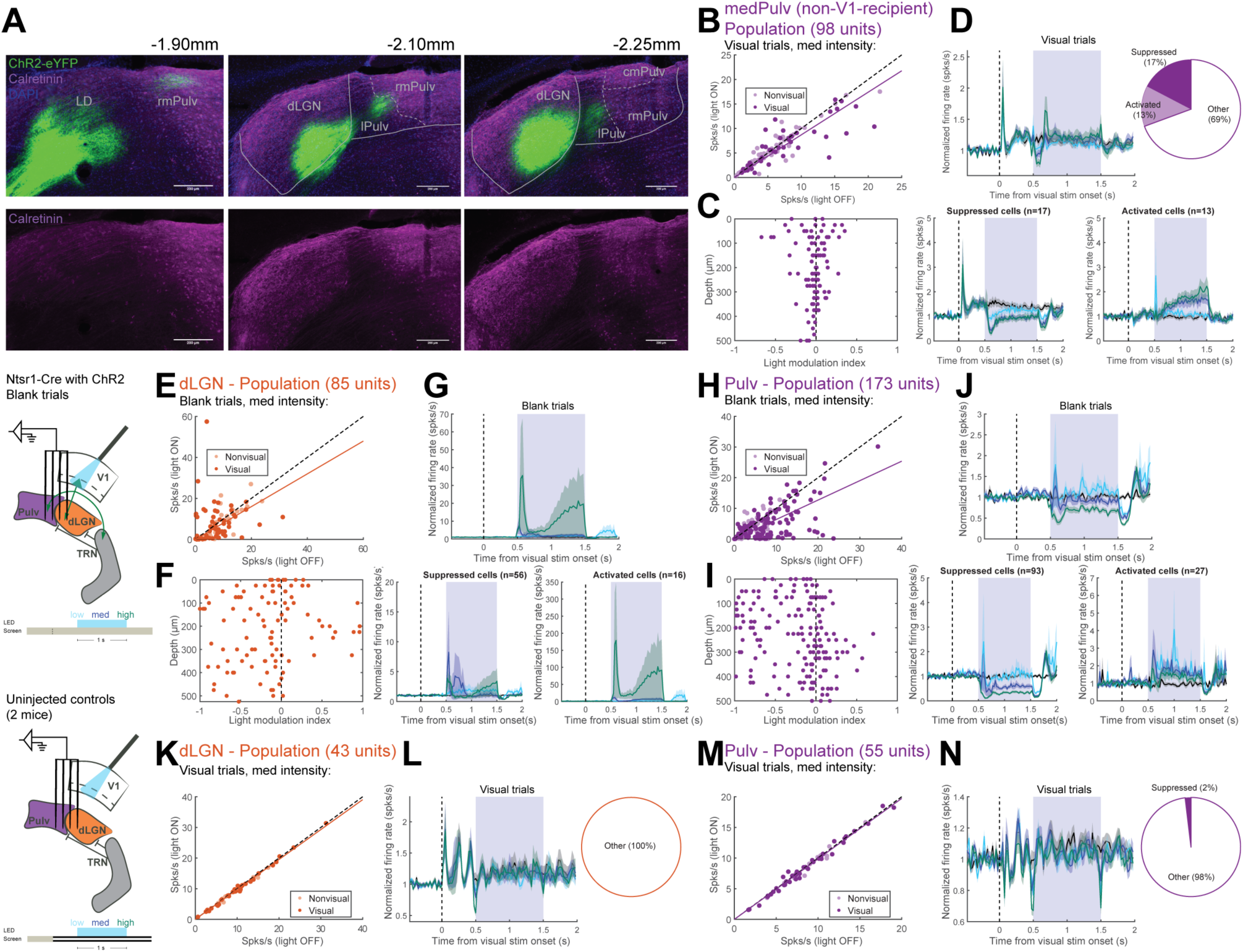
Effects of sustained L6CT photostimulation on different subnuclei of the pulvinar and on spontaneous activity in dLGN and pulvinar. (A) Calretinin helps distinguish between different pulvinar subnuclei. Images in three columns are from different coronal sections in the same animal, from anterior (left) to posterior (right). Bottom row shows the calretinin channel in isolation. (B) Average firing rates during the 1-second photostimulation period from visual trials, with versus without medium-intensity L6CT photostimulation. Saturated points indicate visually-responsive units. (C) Light modulation index by depth (distance from highest channel on the same probe shank and in the same experiment with a visually-responsive unit). (D) Effects of different photostimulation intensities on units’ visually-evoked activity in medPulv. Top, left: average normalized PSTH across all units. Top, right: breakdown of significantly suppressed and activated units recorded in medPulv. Bottom: average normalized PSTH across suppressed (left) and activated (right) units. Shading indicates ±1 standard error of the mean. (E) Average firing rates in dLGN during the 1-second photostimulation period from blank trials with versus without medium-intensity L6CT photostimulation. (F) Light modulation index by depth (calculated from blank trials). (G) Effects of different photostimulation intensities on units’ spontaneous activity in visTRN. Top: average normalized PSTH across all units. Bottom: average normalized PSTH across suppressed (left) and activated (right) units. “Suppressed” and “activated” units are the same as those in Fig. 1G. (H-J) Same as (E-G) but for units recorded in lateral pulvinar. “Suppressed” and “activated” units are the same as those in Fig. 1K. (K) Average firing rates recorded in dLGN from control recordings (no ChR2-eYFP expression) during the 1-second photostimulation period from visual trials, with versus without medium-intensity photostimulation. (L) Left: Average normalized PSTH of all units recorded in dLGN during control recordings. The dip in firing rate at the t and end of the photostimulation period can be attributed to light artifacts at the onset and offset of the LED that impacted accurate spike detection in a narrow time window (within 3ms of LED onset), and so spikes in this window were excluded in all experiments (see Methods). Shading indicates ±1 standard error of the mean. Right: no units were individually significantly modulated by light. (M-N) Same as (K-L) but for units recorded in lateral pulvinar during control experiments. Only one unit was artifactually considered “suppressed” under our criteria for being significantly suppressed or activated.

**Fig. S2.**
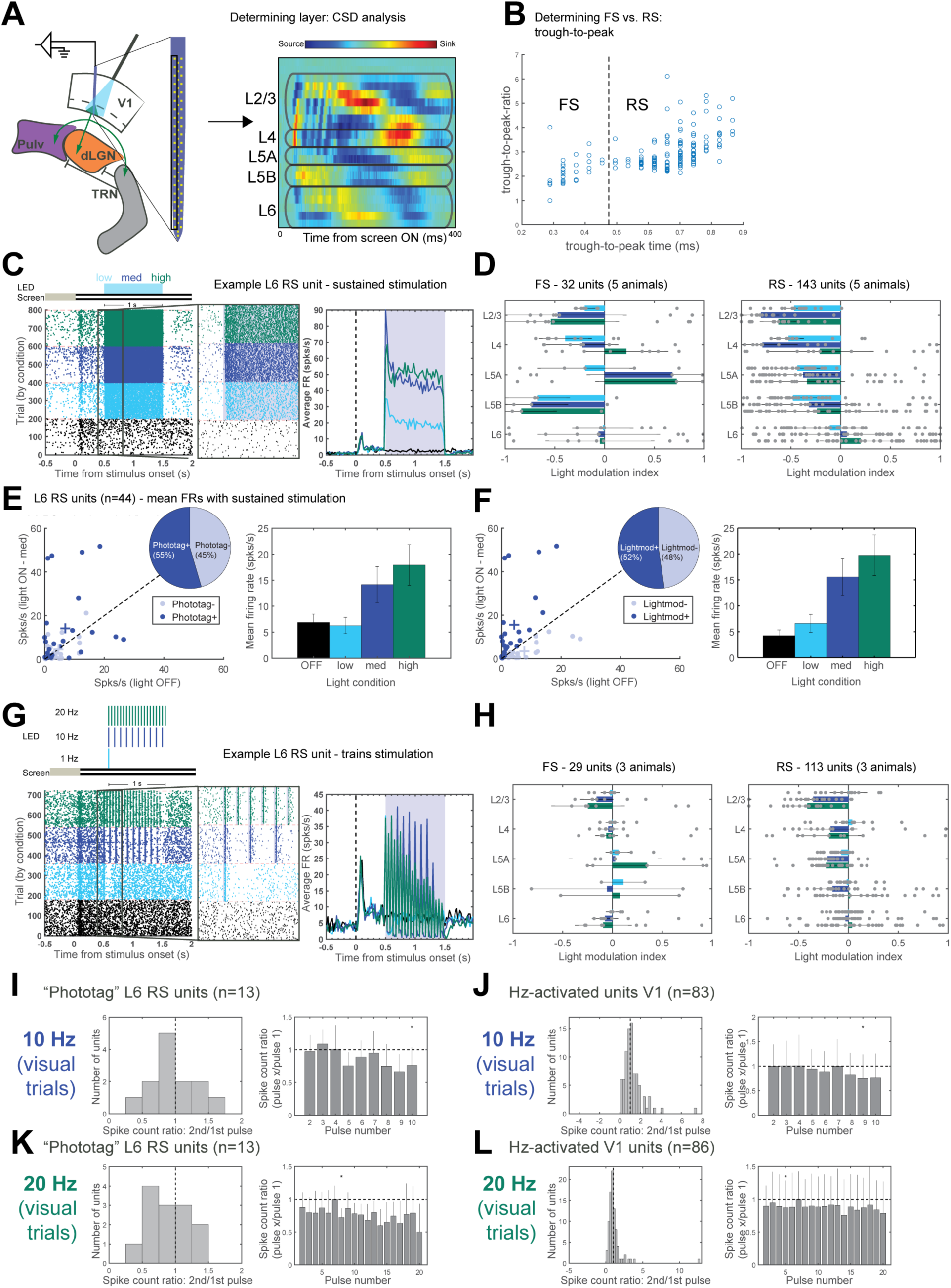
Effects of L6CT optogenetic manipulations on local V1 activity. (A) Schematic of V1 laminar recordings with L6CT photostimulation (left) and determination of which cortical layers correspond to which electrode contacts using current source density (CSD) analysis (right). The CSD plot comes from a single column of electrode contacts on a single probe shank (boxed area). (B) Two distinct groups of units, referred to as fast-spiking (FS) and regular-spiking (RS), are identifiable based on the trough-to-peak times and trough-to-peak ratios of their waveforms. Units with trough-to-peak times less than 0.475ms were considered FS, and others were considered RS. (C) Example L6 RS unit, putatively ChR2-expressing. Left: raster plot organized by photostimulation condition. Middle: zoomed-in image of boxed part of raster plot (from 100ms before to 300ms after LED photostimulation onset), showing short-latency onset of increased activity. Shaded region indicates photostimulation period (1s duration). Right: Average PSTH across visual trials. (D) Median light modulation index by layer of FS (middle) and RS (right) cells with sustained 1-second L6CT photostimulation at low, medium, and high levels. Error bars indicate interquartile range. (E) Left: average firing rates of L6 RS units during the 1-second photostimulation period from visual trials, with and without medium-intensity photostimulation. Saturated points indicate “phototagged”, putative ChR2-expressing L6CT units (see Methods), and crosses indicate mean firing rates across phototagged and non-phototagged units. Inset: proportion of L6 RS units that were phototagged. Right: mean firing rates across all L6 RS “phototagged” units for each light condition. Error bars indicate ±1 standard error of the mean. (F) Same as (E) but with putative ChR2-expressing L6CT units identified by light modulation index > 0. Same example L6 RS unit has in (C) but during the trains experiment, with 1Hz, 10Hz, and 20Hz L6CT photostimulation trains. There was some recoding drift so that more spikes were detected during the trains experiment than in (C). Panel descriptions are same as for (C), although shaded rectangles in left two panels indicate 10ms photostimulation pulses. (H) Like (D) but for trains experiments. (I) Quantification of frequency-dependent spiking of “phototagged” L6 RS units driven at 10Hz. Left: histogram of spike count ratios (comparing spike outputs following the second photostimulation pulse relative to the first) for phototagged units. Right: median spike count ratios across phototagged units, comparing spike outputs following photostimulation pulses 2-10 relative to the first photostimulation pulse. (J) Same as (I) but including all RS and FS units across all layers. (K-L) Same as (I-J) but with 20Hz L6CT photostimulation. Asterisks indicate ratios significantly different from 1 (p’s < 0.05, sign test). Error bars indicate interquartile range.

**Fig. S3.**
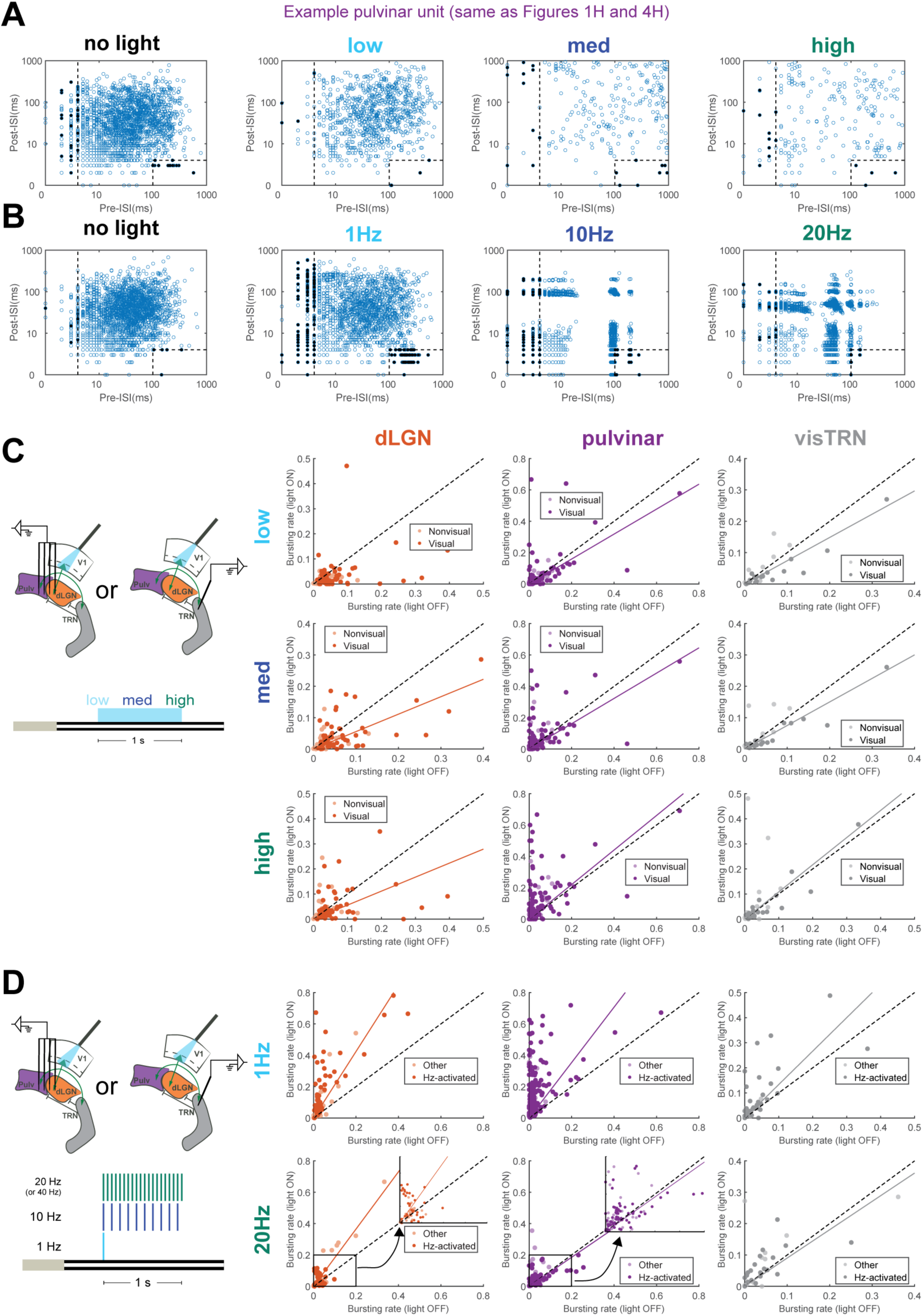
Changes in bursting in dLGN, lateral pulvinar and visTRN under different photostimulation conditions. (A) Scatterplots of pre- and post-interspike intervals (ISIs) for all spikes of an example pulvinar unit (the same as in Fig. 1H, top and Fig. 4H) that occurred during the 1-second photostimulation period during visual trials without photostimulation (left) and during visual trials with low, medium and high continuous L6CT photostimulation (right three panels). Dashed lines demarcate the pre- and post-ISI cut-offs for being considered burst spikes (pre-ISI ≥ 100ms and post-ISI ≤ 4ms, or pre-ISI ≤ 4ms; see Methods). Filled black circles indicate spikes classified as burst spikes (notice that some spikes with preISIs < 4ms were not considered burst spikes if they were not preceded by another burst spike). (B) Same as (A) but for the same unit during train photostimulation experiments with no light, 1 Hz, 10 Hz and 20 Hz photostimulation. (C) Average bursting rates (number of spikes that occurred during bursts / total number of spikes) for all units during visual trials with and without L6CT photostimulation. Left: recording and photostimulation configuration and trial structure. Right: columns are for units recorded in different thalamic nuclei (dLGN, pulvinar, visTRN) and rows are from trials with different degrees of sustained L6CT photostimulation (low, medium, high). (D) Average bursting rates for all units during visual trials with and without L6CT train photostimulation. Left: recording and photostimulation configuration and trial structure. Right: panels as in (C) but during train photostimulation experiments. Plots from 10 Hz photostimulation trials are missing because they are already shown in Figs. 4 and 5.

**Fig. S4.**
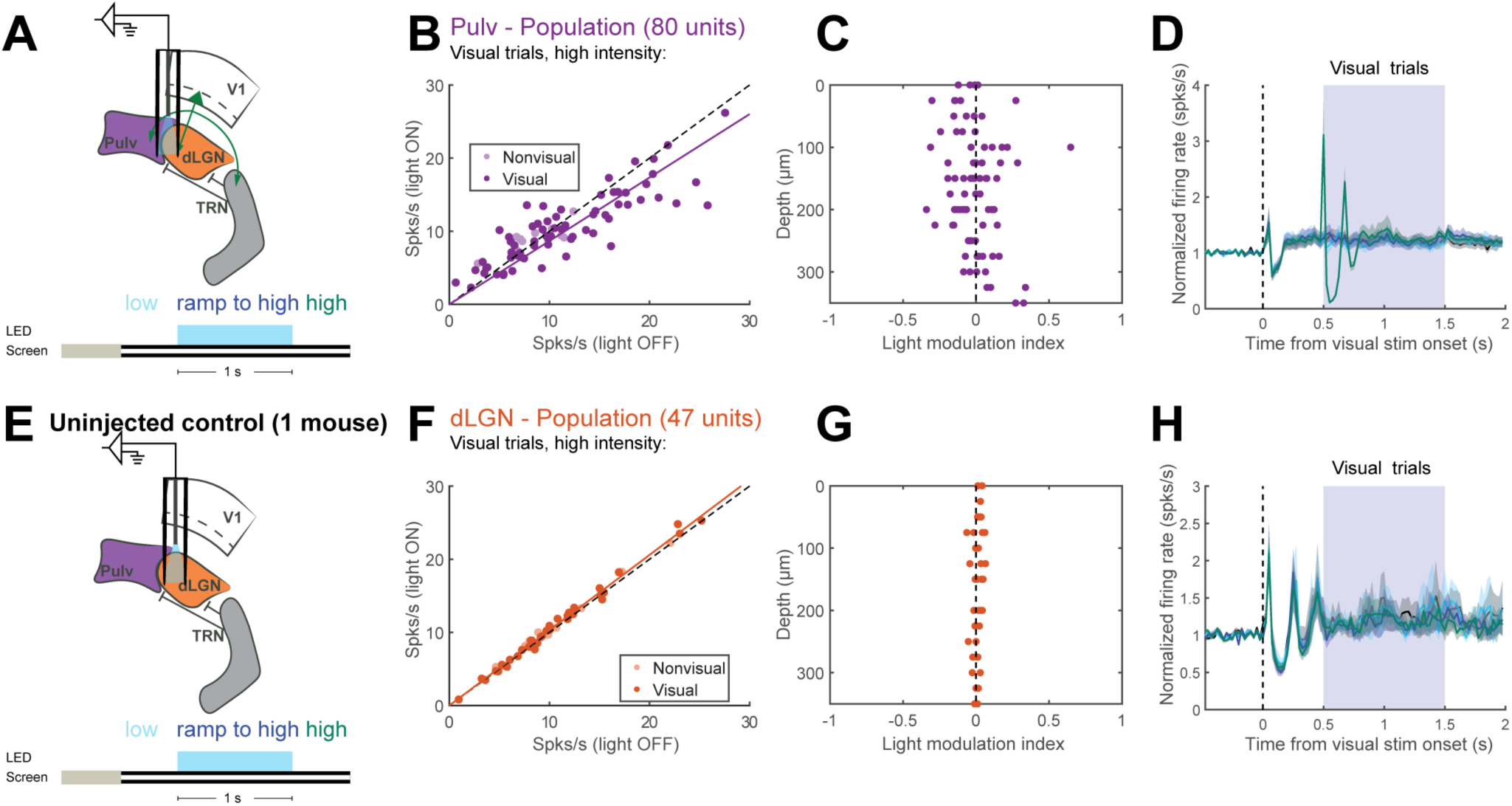
Increased activity with axon terminal stimulation is absent from the pulvinar in experiments using lower light levels and in an uninjected control animal. (A) Diagram of optrode location (half in lateral pulvinar, half in dLGN) and trial structure for visual and LED stimulation. Exact light levels used for “high” photostimulation conditions were titrated per experiment and thus should be interpreted as a relative rather than an absolute value (see Methods). (B) Average firing rates during the 1-second photostimulation period from visual trials, with versus without photostimulation. (C) Light modulation index by depth. (D) Averaged normalized PSTH (visual trials) from all units recorded in the pulvinar during split pulvinar-dLGN optrode recordings. Shading indicates ±1 standard error of the mean. (E) Diagram of optrode location in dLGN and trial structure during a control experiment in an uninjected wild-type mouse. (F) Average firing rates from control experiment during the 1-second photostimulation period from visual trials, with versus without photostimulation. (G) Light modulation index by depth. Average normalized PSTH (visual trials) from all units during the dLGN control optrode recording. Shading indicates ±1 standard error of the mean.

**Fig. S5.**
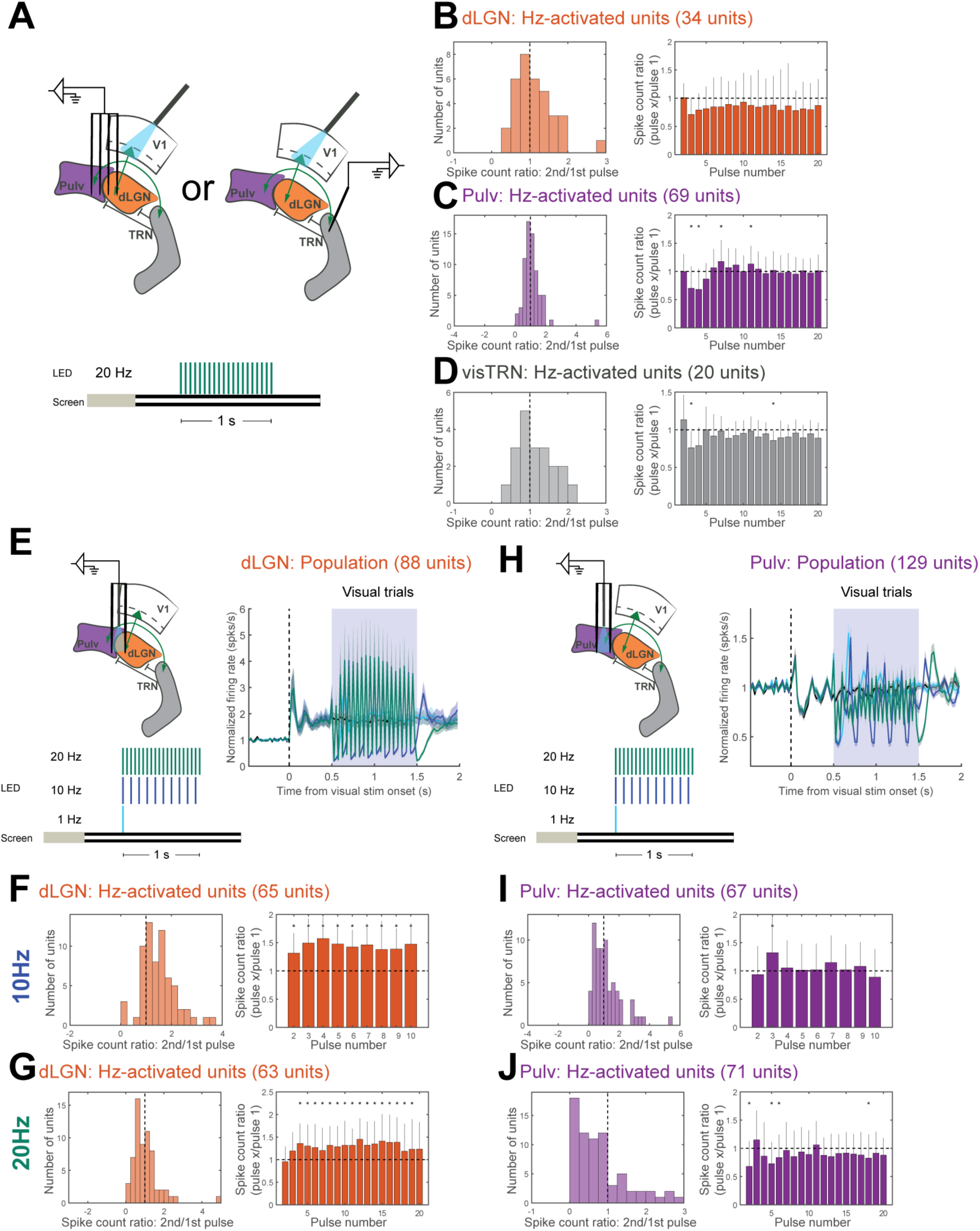
Other frequency-dependent effects using 20Hz cell body photostimulation and axon terminal stimulation. (A) Diagram of recording and photostimulation configuration in dLGN/pulvinar and visTRN recordings and trial structure of visual trials with 20Hz L6CT photostimulation. Experiments from two animals in dLGN/pulvinar that were included in the 10Hz photostimulation analysis in Fig. 4 were excluded from these analyses because the high-frequency condition was 40Hz instead of 20Hz. (B) Quantification of dLGN spiking with 20Hz L6CT photostimulation. Left: histogram of spike count ratios (comparing spike outputs following the second photostimulation pulse relative to the first) for 20Hz-activated units. Right: median spike count ratios across Hz-activated units, comparing spike outputs following photostimulation pulses 2-20 relative to the first photostimulation pulse. Asterisks indicate ratios significantly different from 1 (p’s < 0.05, sign test). Error bars indicate interquartile range. (C) Quantification of pulvinar spiking with 20Hz L6CT photostimulation. Panel descriptions same as for (B). (D) Quantification of visTRN spiking with 20Hz L6CT photostimulation. Panel descriptions same as for (B). (E) Photostimulation trains experiments during dLGN optrode recordings. Left: diagram of optrode configuration (partially or entirely in dLGN) and trial structure for visual and LED stimulation during trains experiments. Right: average normalized PSTH of dLGN units’ activity during visual trials. Shading indicates ±1 standard error of the mean. (F) Quantification of dLGN spiking with 10Hz axon terminal stimulation. Panel descriptions same as for (B). (G) Quantification of dLGN spiking with 20Hz axon terminal stimulation. Panel descriptions as for (B). (H) Photostimulation trains experiments during pulvinar optrode recordings. Left: diagram of optrode configuration in the pulvinar and trial structure for visual and LED stimulation during trains experiments. Right: average normalized PSTH of pulvinar units’ activity during visual trials. Shading indicates ±1 standard error of the mean. (I-J) Quantification of pulvinar spiking with 10Hz (I) and 20Hz (J) axon terminal stimulation. Panel descriptions same as for (B).

## References

1. Sherman SM, Guillery RW (1998) On the actions that one nerve cell can have on another: distinguishing “drivers” from “modulators”. Proc Natl Acad Sci USA 95(12):7121–7126.

2. Przybyszewski AW, Gaska JP, Foote W, Pollen DA (2000) Striate cortex increases contrast gain of macaque LGN neurons. Vis Neurosci 17(4):485–494.

3. Olsen SR, Bortone DS, Adesnik H, Scanziani M (2012) Gain control by layer six in cortical circuits of vision. Nature 483(7387):47–52.

4. Denman DJ, Contreras D (2015) Complex Effects on In Vivo Visual Responses by Specific Projections from Mouse Cortical Layer 6 to Dorsal Lateral Geniculate Nucleus. J Neurosci 35(25):9265–9280.

5. Wang W, Andolina IM, Lu Y, Jones HE, Sillito AM (2018) Focal gain control of thalamic visual receptive fields by layer 6 corticothalamic feedback. Cereb Cortex 28(1):267–280.

6. Wang W, Jones HE, Andolina IM, Salt TE, Sillito AM (2006) Functional alignment of feedback effects from visual cortex to thalamus. Nat Neurosci 9(10):1330–1336.

7. Andolina IM, Jones HE, Wang W, Sillito AM (2007) Corticothalamic feedback enhances stimulus response precision in the visual system. Proc Natl Acad Sci USA 104(5):1685–1690.

8. Hasse JM, Briggs F (2017) Corticogeniculate feedback sharpens the temporal precision and spatial resolution of visual signals in the ferret. Proc Natl Acad Sci USA 114(30):E6222–E6230.

9. Andolina IM, Jones HE, Sillito AM (2013) Effects of cortical feedback on the spatial properties of relay cells in the lateral geniculate nucleus. J Neurophysiol 109(3):889–899.

10. Mease RA, Krieger P, Groh A (2014) Cortical control of adaptation and sensory relay mode in the thalamus. Proc Natl Acad Sci USA 111(18):6798–6803.

11. Bourassa J, Deschênes M (1995) Corticothalamic projections from the primary visual cortex in rats: a single fiber study using biocytin as an anterograde tracer. Neuroscience 66(2):253–263.

12. Li J, Wang S, Bickford ME (2003) Comparison of the ultrastructure of cortical and retinal terminals in the rat dorsal lateral geniculate and lateral posterior nuclei. J Comp Neurol 460(3):394–409.

13. Li J, Guido W, Bickford ME (2003) Two distinct types of corticothalamic EPSPs and their contribution to short-term synaptic plasticity. J Neurophysiol 90(5):3429–3440.

14. Reichova I, Sherman SM (2004) Somatosensory corticothalamic projections: distinguishing drivers from modulators. J Neurophysiol 92(4):2185–2197.

15. Guo W, Clause AR, Barth-Maron A, Polley DB (2017) A Corticothalamic Circuit for Dynamic Switching between Feature Detection and Discrimination. Neuron 95(1):180–194.e5.

16. Pauzin FP, Krieger P (2018) A corticothalamic circuit for refining tactile encoding. Cell Rep 23(5):1314–1325.

17. Durand S, et al. (2016) A comparison of visual response properties in the lateral geniculate nucleus and primary visual cortex of awake and anesthetized mice. J Neurosci 36(48):12144–12156.

18. Crandall SR, Cruikshank SJ, Connors BW (2015) A corticothalamic switch: controlling the thalamus with dynamic synapses. Neuron 86(3):768–782.

19. Gong S, et al. (2007) Targeting Cre recombinase to specific neuron populations with bacterial artificial chromosome constructs. J Neurosci 27(37):9817–9823.

20. Bortone DS, Olsen SR, Scanziani M (2014) Translaminar inhibitory cells recruited by layer 6 corticothalamic neurons suppress visual cortex. Neuron 82(2):474–485.

21. Yang L, Lee K, Villagracia J, Masmanidis SC (2019) Open source silicon microprobes for high throughput neural recording. J Neural Eng.

22. Beltramo R, Scanziani M (2019) A collicular visual cortex: Neocortical space for an ancient midbrain visual structure. Science 363(6422):64–69.

23. Zhou NA, Maire PS, Masterson SP, Bickford ME (2017) The mouse pulvinar nucleus: Organization of the tectorecipient zones. Vis Neurosci 34:E011.

24. Bennett C, et al. (2019) Higher-Order Thalamic Circuits Channel Parallel Streams of Visual Information in Mice. Neuron 102(2):477–492.e5.

25. Pinault D, Deschênes M (1998) Projection and innervation patterns of individual thalamic reticular axons in the thalamus of the adult rat: a three-dimensional, graphic, and morphometric analysis. J Comp Neurol 391(2):180–203.

26. Jurgens CWD, Bell KA, McQuiston AR, Guido W (2012) Optogenetic stimulation of the corticothalamic pathway affects relay cells and GABAergic neurons differently in the mouse visual thalamus. PLoS One 7(9):e45717.

27. Cruikshank SJ, Urabe H, Nurmikko AV, Connors BW (2010) Pathway-specific feedforward circuits between thalamus and neocortex revealed by selective optical stimulation of axons. Neuron 65(2):230–245.

28. Williamson RS, Polley DB (2019) Parallel pathways for sound processing and functional connectivity among layer 5 and 6 auditory corticofugal neurons. Elife 8.

29. Mattis J, et al. (2011) Principles for applying optogenetic tools derived from direct comparative analysis of microbial opsins. Nat Methods 9(2):159–172.

30. Hass CA, Glickfeld LL (2016) High-fidelity optical excitation of cortico-cortical projections at physiological frequencies. J Neurophysiol 116(5):2056–2066.

31. Briggs F, Usrey WM (2008) Emerging views of corticothalamic function. Curr Opin Neurobiol 18(4):403–407.

32. McCormick DA, von Krosigk M (1992) Corticothalamic activation modulates thalamic firing through glutamate “metabotropic” receptors. Proc Natl Acad Sci USA 89(7):2774–2778.

33. Tsumoto T, Creutzfeldt OD, Legéndy CR (1978) Functional organization of the corticofugal system from visual cortex to lateral geniculate nucleus in the cat (with an appendix on geniculo-cortical mono-synaptic connections). Exp Brain Res 32(3):345–364.

34. Li L, Ebner FF (2007) Cortical modulation of spatial and angular tuning maps in the rat thalamus. J Neurosci 27(1):167–179.

35. Conley M, Diamond IT (1990) Organization of the visual sector of the thalamic reticular nucleus in galago. Eur J Neurosci 2(3):211–226.

36. Jones EG ed. (1985) The Thalamus (Plenum Press, New York).

37. Evangelio M, García-Amado M, Clascá F (2018) Thalamocortical projection neuron and interneuron numbers in the visual thalamic nuclei of the adult C57BL/6 mouse. Front Neuroanat 12:27.

38. Augustinaite S, Yanagawa Y, Heggelund P (2011) Cortical feedback regulation of input to visual cortex: role of intrageniculate interneurons. J Physiol (Lond) 589(Pt 12):2963–2977.

39. Granseth B (2004) Dynamic properties of corticogeniculate excitatory transmission in the rat dorsal lateral geniculate nucleus in vitro. J Physiol (Lond) 556(Pt 1):135–146.

40. von Krosigk M, Monckton JE, Reiner PB, McCormick DA (1999) Dynamic properties of corticothalamic excitatory postsynaptic potentials and thalamic reticular inhibitory postsynaptic potentials in thalamocortical neurons of the guineapig dorsal lateral geniculate nucleus. Neuroscience 91(1):7–20.

41. Lindström S, Wróbel A (1990) Frequency dependent corticofugal excitation of principal cells in the cat’s dorsal lateral geniculate nucleus. Exp Brain Res 79(2):313–318.

42. Jahnsen H, Llinás R (1984) Electrophysiological properties of guinea-pig thalamic neurones: an in vitro study. J Physiol (Lond) 349:205–226.

43. Jahnsen H, Llinás R (1984) Ionic basis for the electro-responsiveness and oscillatory properties of guinea-pig thalamic neurones in vitro. J Physiol (Lond) 349:227–247.

44. Anderson MP, et al. (2005) Thalamic Cav3.1 T-type Ca2+ channel plays a crucial role in stabilizing sleep. Proc Natl Acad Sci USA 102(5):1743–1748.

45. Sherman SM (2016) Thalamus plays a central role in ongoing cortical functioning. Nat Neurosci 19(4):533–541.

46. Roth MM, et al. (2016) Thalamic nuclei convey diverse contextual information to layer 1 of visual cortex. Nat Neurosci 19(2):299–307.

47. Chevée M, Robertson JDJ, Cannon GH, Brown SP, Goff LA (2018) Variation in activity state, axonal projection, and position define the transcriptional identity of individual neocortical projection neurons. Cell Rep 22(2):441–455.

48. Bourassa J, Pinault D, Deschênes M (1995) Corticothalamic projections from the cortical barrel field to the somatosensory thalamus in rats: a single-fibre study using biocytin as an anterograde tracer. Eur J Neurosci 7(1):19–30.

49. Hoerder-Suabedissen A, et al. (2018) Subset of cortical layer 6b neurons selectively innervates higher order thalamic nuclei in mice. Cereb Cortex 28(5):1882–1897.

50. Groh A, et al. (2010) Cell-type specific properties of pyramidal neurons in neocortex underlying a layout that is modifiable depending on the cortical area. Cereb Cortex 20(4):826–836.

51. Jaramillo J, Mejias JF, Wang X-J (2019) Engagement of Pulvino-cortical Feedforward and Feedback Pathways in Cognitive Computations. Neuron 101(2):321–336.e9.

52. Siegle JH, et al. (2017) Open Ephys: an open-source, plugin-based platform for multichannel electrophysiology. J Neural Eng 14(4):045003.

53. Marshel JH, Garrett ME, Nauhaus I, Callaway EM (2011) Functional specialization of seven mouse visual cortical areas. Neuron 72(6):1040–1054.

54. Pachitariu M, Steinmetz N, Kadir S, Carandini M, Harris KD (2016) Kilosort: realtime spike-sorting for extracellular electrophysiology with hundreds of channels. BioRxiv.

55. Rossant C, et al. (2016) Spike sorting for large, dense electrode arrays. Nat Neurosci 19(4):634–641.

56. Schmitzer-Torbert N, Jackson J, Henze D, Harris K, Redish AD (2005) Quantitative measures of cluster quality for use in extracellular recordings. Neuroscience 131(1):1–11.

57. Niell CM, Stryker MP (2010) Modulation of visual responses by behavioral state in mouse visual cortex. Neuron 65(4):472–479.

58. Erisken S, et al. (2014) Effects of locomotion extend throughout the mouse early visual system. Curr Biol 24(24):2899–2907.

59. Reinagel P, Godwin D, Sherman SM, Koch C (1999) Encoding of visual information by LGN bursts. J Neurophysiol 81(5):2558–2569.

## SI References

1. Pettersen KH, Devor A, Ulbert I, Dale AM, Einevoll GT (2006) Current-source density estimation based on inversion of electrostatic forward solution: effects of finite extent of neuronal activity and conductivity discontinuities. J Neurosci Methods 154(1–2):116–133.

2. Kvitsiani D, et al. (2013) Distinct behavioural and network correlates of two interneuron types in prefrontal cortex. Nature 498(7454):363–366.

